# Structures of BIRC6-Client Complexes Provide Mechanism of Smac-Mediated Release of Caspases

**DOI:** 10.1101/2022.08.19.504532

**Authors:** Moritz Hunkeler, Cyrus Y. Jin, Eric S. Fischer

## Abstract

Apoptosis is tightly regulated and essential for metazoan development^1, 2^. Excessive apoptosis contributes to neurodegenerative disease, while diminished apoptosis can lead to inflammation and cancer^3^. Inhibitor of apoptosis (IAP) proteins are the principal actors that restrain apoptotic activity and are thus attractive therapeutic targets^4^. IAPs in turn are regulated by mitochondria-derived pro-apoptotic factors such as Smac and HtrA2^4^. Here, through a series of cryo-electron microscopy (cryo-EM) structures of full-length baculoviral IAP repeat-containing protein 6 (BIRC6) bound to Smac, caspase-3, caspase-7 and HtrA2, we provide the molecular basis for BIRC6-mediated caspase inhibition and its release by Smac. We demonstrate that BIRC6 cooperates preferentially with the non-canonical E1 enzyme UBA6 to ubiquitylate caspases and that caspase ubiquitylation is effectively inhibited by Smac through near-irreversible interactions. The dimeric arrangement of BIRC6 resolves the long-standing question of how the evolutionarily conserved single-BIR domain IAPs sequester caspases and how Smac binding antagonizes IAPs. This collection of BIRC6 structures provides critical insights into IAP-mediated apoptosis regulation, relevant for the development of current and future apoptosis-targeting cancer therapeutics.

## Introduction

Genetically encoded cell death programs such as apoptosis, necroptosis or pyroptosis remove infected, damaged or obsolete cells during development and are essential in all metazoans^1^. Aberrant activity or lack of regulation of cell death pathways leads to a wide array of human pathologies such as neurodegeneration, auto-inflammatory disease and cancer^2, 3^. Apoptosis, unlike necroptosis and pyroptosis, results in phagocyte-dependent cell death without leakage of cellular contents or proinflammatory molecules^2^. Extrinsic, initiated by external death signals transduced through death receptors, and intrinsic, initiated by internal cues such as excessive DNA or mitochondrial damage^2^, apoptotic signaling pathways both use activation of cysteine-dependent aspartyl specific proteases (caspases) to irreversibly cleave a plethora of cellular substrates and trigger execution of the apoptotic program^5^.

Inhibitor of apoptosis (IAP) proteins are the principal anti-apoptotic factors that keep caspases and cell death commitment in check by direct binding and inhibition of initiator caspase-9 (casp- 9) and executioner caspases-3 and 7 (casp-3 and casp-7)^4^. Many cancer cells express elevated levels of IAPs and have heightened apoptotic thresholds to resist apoptotic signals and cytotoxic therapies^6^. In mammals, seven IAPs, also referred to as BIRCs (BIR domain-containing proteins) exist, which have been classified into two classes. Class I contains BIRC1/NAIP, BIRC2/cIAP-1, BIRC3/cIAP-2, BIRC4/XIAP and BIRC7/ML-IAP, while class II is comprised of BIRC5/survivin and BIRC6/Apollon^4, 7, 8^. The BIR domain found in all IAPs interacts with the conserved IAP binding motif (IBM) of caspases^9^. Similar IBMs are found on Smac and HtrA2 upon release from the mitochondria^9^. Anti-apoptotic IAPs (all except for survivin) have acquired ubiquitin E3 ligase activity necessary for anti-apoptotic activity^4, 7^. Intrinsic and extrinsic apoptotic stimuli, if accumulating, ultimately lead to BCL-2 family-mediated release of pro-apoptotic molecules from the mitochondria including cytochrome c^10^, apoptosis inducing factor^11^, second mitochondrial activator of caspases (Smac, also known as direct IAP-binding protein with low isoelectric point (DIABLO))^12, 13^, and the serine protease High temperature requirement protein A2 (HtrA2)^14^. Smac and HtrA2 both directly interact with IAPs and block IAP-mediated caspase inhibition thereby releasing the brakes on cell death^4^. These findings lead to the development of multiple small molecule Smac-mimetics that target the IAP/caspase interaction to promote apoptosis as anti-cancer therapeutics^15^, several currently undergoing clinical exploration^16^. Despite these translational efforts, how IAPs and specifically the class II IAP BIRC6 recognize caspases and how this is antagonized by Smac remains poorly understood. This lack of understanding prevents the development of more effective drugs targeting this important node of apoptosis.

BIRC6 (also called Apollon^17^ or Bruce^18^) plays critical roles in cell division^19, 20^, and was shown to exhibit prototypical anti-apoptotic activity by directly antagonizing caspase activity^19, 21^. BIRC6 is the only essential IAP^21^ and is evolutionarily the oldest IAP to acquire E3 ubiquitin ligase activity^22^. The large, multidomain 530-kDa protein has a single BIR domain close to the amino-terminus (N-terminus) and an E2-E3 hybrid ubiquitin-conjugating (UBC) domain close to the carboxy-terminus (C-terminus), while the rest of the protein remains largely uncharacterized (**Fig. 1a**). BIRC6 interacts with and inhibits casp-3, casp-7 and casp-9, and these interactions are inhibited by Smac^19^. BIRC6 has also been established as a substrate for caspases as well as HtrA2 (**Fig. 1b**)^19^. In addition to acting as an IAP and playing crucial roles in orchestrating the progression through cytokinesis^20, 23^, BIRC6 was found to regulate autophagy^24, 25^, thereby connecting key cellular pathways and apoptotic control.

**Fig. 1.**
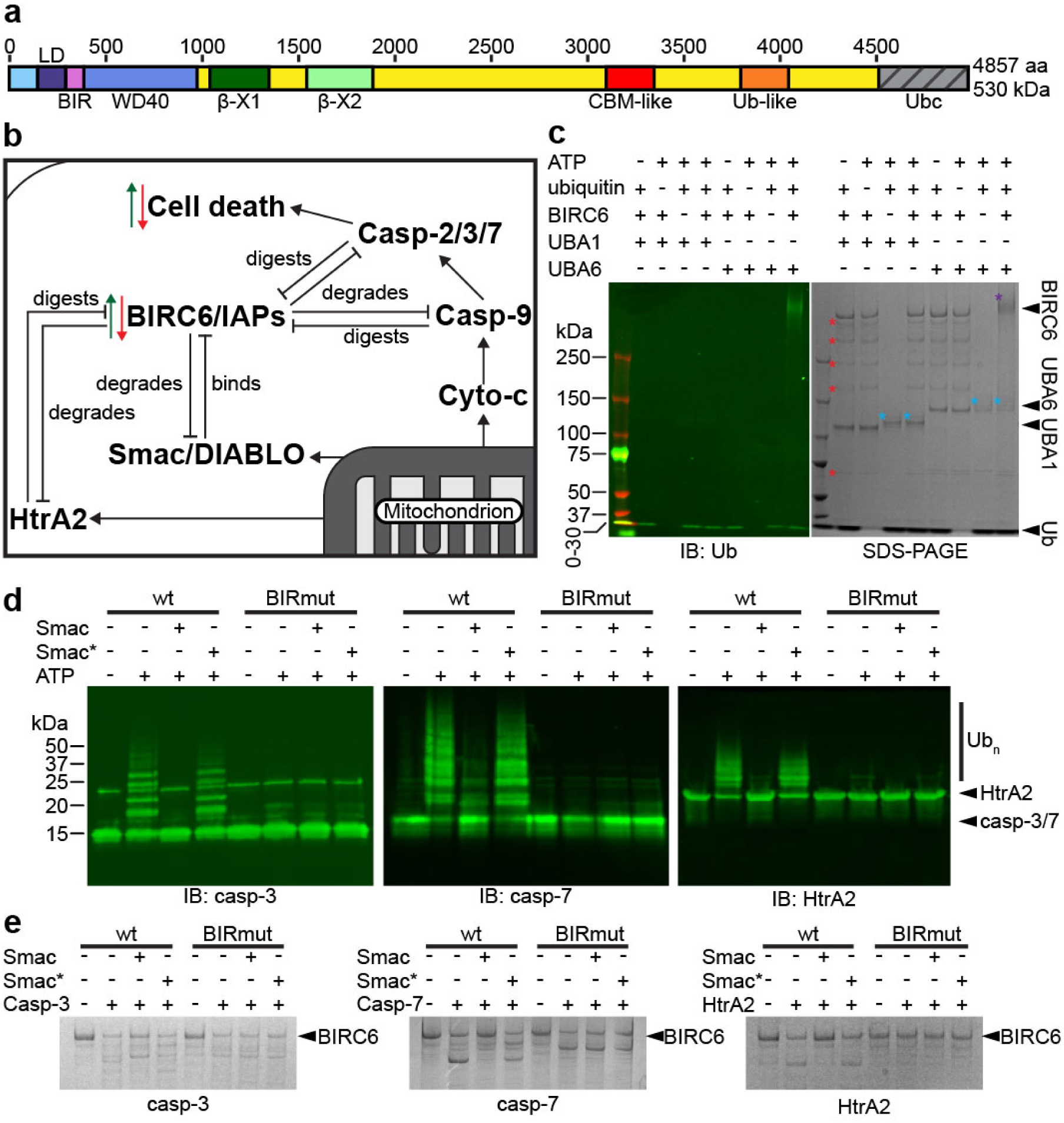
BIRC6 is a UBA6-dependent ubiquitin ligase for caspases. **a**, Domain scheme for BIRC6. Domain coloring and labeling is held constant throughout the manuscript. **b**, Schematic overview of reported cellular interactions of BIRC6. **c**, Auto-ubiquitylation assay. Anti-ubiquitin western blot (left) and SDS-PAGE (right) analysis of BIRC6 auto-ubiquitylation. Purple asterisk indicates auto-ubiquitylated BIRC6, red asterisks are BIRC6 degradation bands. Blue asterisks indicate ubiquitin-charged E1. **d**, Ubiquitylation assay. Western blots using the indicated antibodies of ubiquitylation assays establishing casp-3, casp-7 and HtrA2 as in vitro substrates of BIRC6. Addition of wild-type processed Smac (N-terminal AVPI) inhibits activity, but not addition of a Smac variant (Smac*, N-terminal MVPI). **e**, SDS-PAGE analysis of BIRC6 stability assays upon incubation with casp-3, casp-7 and HtrA2. All three proteases digest wild-type (wt) BIRC6 (black arrow), and the effect can be reverted to base line degradation by adding Smac (but not Smac*). Base line degradation of a BIR domain mutant (BIRmut, C328S/C331S) is insensitive to the presence of Smac or Smac*. All gels and blots are representative for at least three independent replicates.

To date, only structures of isolated domains or fragments have been reported for IAPs and IAP-containing complexes. Due to this lack of structural information, it remains elusive how IAPs and specifically BIRC6, bind to and inhibit pro-apoptotic caspases and how this is effectively counteracted by Smac and HtrA2. We have biochemically characterized and obtained several cryo-electron microscopy (cryo-EM) structures of full-length BIRC6 alone or bound to Smac, casp-3, casp-7 or HtrA2. These establish a molecular understanding of how IAPs engage diverse substrates and inhibitors to control apoptosis. We find that an unexpected dimeric architecture of BIRC6 establishes an accommodating central cavity that allows for competitive binding of diverse factors recruited through the BIR domain and stabilized by electrostatic interactions. This architecture, together with near irreversible binding of Smac, reconciles the long-standing question of how caspase inhibition by IAPs is released upon apoptosis.

## Results

### BIRC6 is a UBA6-dependent E2/E3 chimera

BIRC6 has been demonstrated to exhibit UBC domain-dependent anti-apoptotic activity and to bind to caspases, HtrA2 and Smac through its BIR domain in cells^19, 20, 23–25^. To reconstitute these activities in a fully recombinant system and enable structural studies, we expressed and purified full-length BIRC6 (**Extended Data Fig. 1a**). By examining the Cancer Dependency Map (DepMap^26^), we noticed a strong correlation between BIRC6 and the non-canonical ubiquitin activating enzyme UBA6. Based on this observation we set out to test whether BIRC6 prefers UBA6 over the canonical ubiquitin E1 enzyme UBA1. Auto-ubiquitylation assays using UBA1 or UBA6 established a clear preference for UBA6 with negligible activity observed with UBA1 (**Fig. 1c**), consistent with previous observations that UBA6 and BIRC6 cooperate to regulate autophagy^25^. We further confirmed that UBA6 was also the preferred E1 for BIRC6-dependent ubiquitylation of casp-3, casp-7 and HtrA2 (**Extended Data Fig. 1b**). BIRC6 incubated with inhibited casp-3, casp-7 or HtrA2-S306A in presence of UBA6, ubiquitin and ATP resulted in efficient ubiquitylation of all three proteases, which is strictly dependent on the BIR domain (**Fig. 1d**). The ubiquitylation of all substrates is inhibited by Smac (**Fig. 1d**), which itself is also a substrate for BIRC6 ubiquitylation in vitro^27^ (**Extended Data Fig. 1c,d**). A mutant form of Smac (Smac*, IBM motif mutated to MVPI), which no longer binds to BIR domains^19^, fails to inhibit BIRC6 activity (**Fig. 1d**). In absence of protease inhibitors, casp-3, casp-7 and HtrA2 digest BIRC6 in a strictly BIR domain-dependent fashion, which is inhibited by Smac but not Smac* (**Fig. 1e**). Together, these findings establish that BIRC6 acts as an IAP utilizing its BIR and UBC domains to bind and ubiquitylate casp-3, casp-7 and HtrA2 and that BIRC6 itself is a substrate for these proteases. The absence of multiple BIR domains and linker regions previously shown to be important for efficient caspase binding^28^ raises the question of how BIRC6 interacts with caspases and its regulators Smac and HtrA2.

### Cryo-EM structure reveals horseshoe-shaped dimeric architecture

To structurally characterize BIRC6 we collected cryo-EM data of full-length BIRC6. Initial two-dimensional (2D) classification revealed the presence of high-quality particles but also significant preferred orientations leading to highly anisotropic three-dimensional (3D) reconstructions (**Extended Data Fig. 2**). Omitting 2D classification and directly employing 3D classification allowed retention of particles representing rare views, which mitigated the preferred orientation problem. Several rounds of classification and local refinements lead to high-quality maps of BIRC6 refined to resolutions of 2.0-3.0 Å (**Fig. 2a and Extended Data Fig. 2 and 3**). BIRC6 presents itself as a large (180 x 170 x 120 Å) head-to-tail dimer with an extended helical arch forming the dimer interface (∼10,286 Å^2^ interface area) and serving as the backbone for functional domains (**Fig. 2b**). At the very N-terminus, we find an unpredicted WD40-like propeller domain, which has a linker domain (LD) and the BIR domain protruding out between individual blades of the propeller (**Fig. 2c**). The IBM binding groove on the BIR domain is facing the central cavity of BIRC6, as revealed by superposition of crystal structures of the XIAP BIR domain in complex with the AVPI peptide of Smac^29^ (**Fig. 2c**). The three N-terminal domains are connected to the helical arch by a disordered linker (∼40 amino acids (aa)) and are sitting on two beta-sandwich domains (β-X1 and β-X2). Centrally inserted to the helical arch are two carbohydrate binding module family 32 (CBM)-like domains^30^ (**Fig. 2b,d**), followed by an unpredicted ubiquitin like domain (Ubl) close to the C-terminal UBC domain (**Fig. 2e**). The UBC domain itself is invisible in the cryo-EM reconstructions due to its high degree of flexibility, which likely enables efficient ubiquitylation of diverse substrates.

**Fig. 2.**
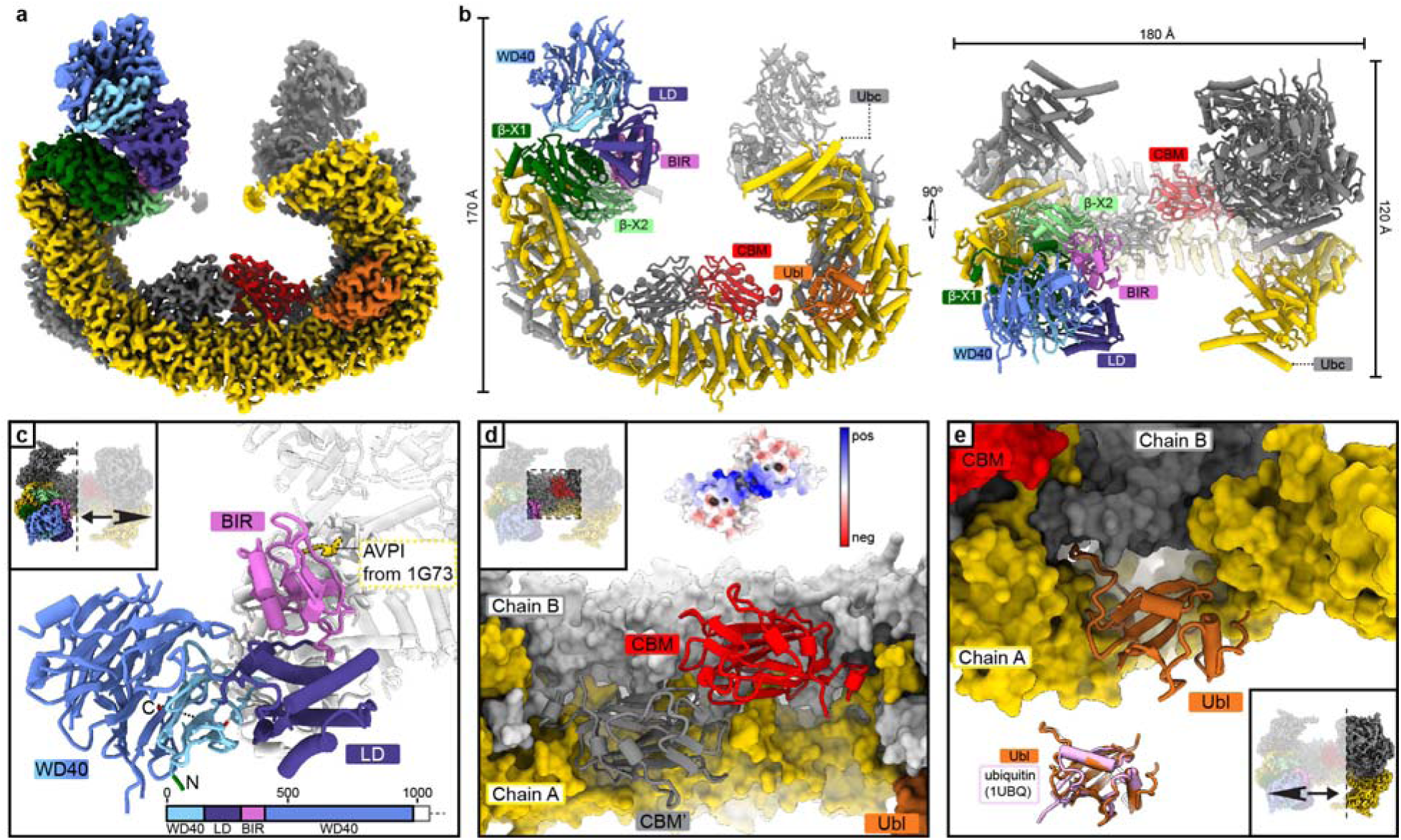
Cryo-EM structure of full-length human BIRC6. **a**, Composite EM map of the BIRC6 dimer, with one chain colored as in Fig. 1a and the other chain colored in grey. **b**, Cartoon model of BIRC6 in two orientations. Domains of one protomer are labeled. The UBC domain is not visible in the reconstructions. **c**, The N-terminal ∼1000 aa comprise a disconnected WD40 propeller with the LD and BIR domains inserted between blades 2 and 3. The location of the peptide binding grove on the BIR domain is illustrated by placement of the AVPI peptide taken from PDB 1G73. The N- and C-termini are colored green and red, respectively, and the disordered linker between the C-terminus and the β-X1 domain is indicated by a dashed line behind the domains. **d**, Close up on the CBM-like domains, holding the dimer together in a ball clasp fashion. The coloumb surface of the two domains is shown on the top, revealing a highly positively charged path right in the center. **e**, Zoomed-in view of the unpredicted ubiquitin-like domain (Ubl). An overlay with ubiquitin (1UBQ) is shown at the bottom.

### Tight binding of Smac to BIRC6 facilitates caspase release mechanism

Previous reports had found similar affinities for Smac and caspase binding to XIAP and cIAP^31, 32^ at odds with effective release of caspase inhibition by Smac. To reconcile these observations, a multivalent binding model with two distinct BIR domains tucked under an arch-shaped Smac dimer was proposed^29, 32, 33^. This model cannot be universal for all IAPs as BIRC6 only has a single BIR domain. In addition, even if one BIR domain was contributed from each protomer, they are still too far apart from each other to form the proposed arrangement (**Extended Data Fig. 4a**). We thus set out to characterize the binding mode of Smac to BIRC6 and established a time-resolved Förster resonance energy transfer (TR-FRET)-based equilibrium binding assay to determine binding constants of Smac, casp-3, casp-7 and HtrA2-S306A. Smac binding is the tightest observed, with an apparent *K*_D_ below 2 nM (assay limited), followed by casp-7 and HtrA2-S306A with 4 ± 1 nM (assay limited) and 75 ± 5 nM, respectively (**Extended Data Fig. 4b,c**). Binding of casp-3 was too weak to establish a *K*_D_. To determine the rank order of competitive binding relevant for regulation, we established a TR-FRET displacement assay where BODIPY-labeled HtrA2-S306A was displaced from BIRC6 by titration of unlabeled Smac, casp-3, casp-7 and HtrA-S306A with IC_50_ values of 9 ± 1 nM, 167 ± 31 nM, 383 ± 60 nM and 1358 ± 148 nM for Smac, casp-7, HtrA2 and casp-3, respectively (**Fig. 3a**). These affinities are consistent with mature Smac released from the mitochondria effectively inhibiting the anti-apoptotic activity of BIRC6 by blocking interactions with caspases.

**Fig. 3.**
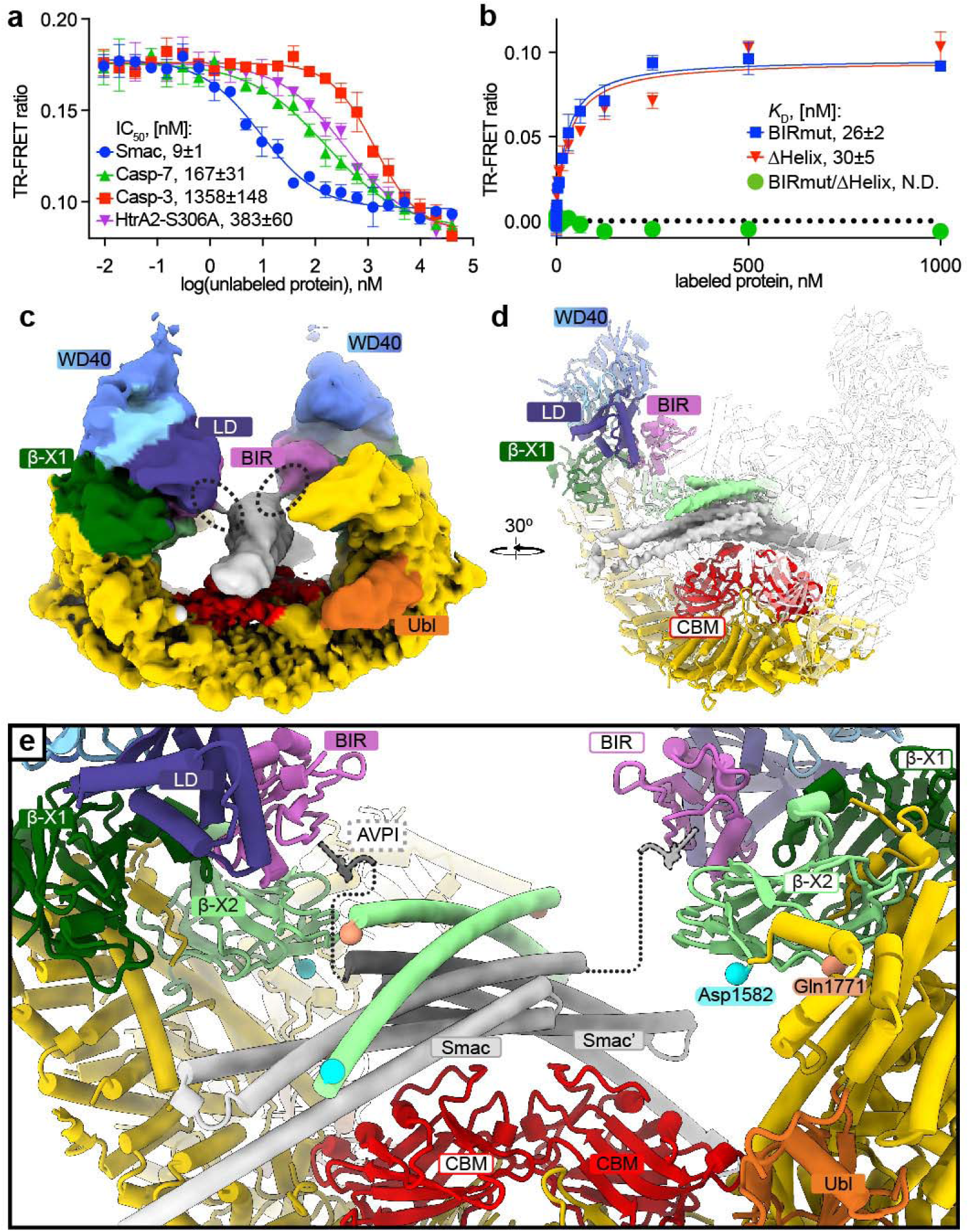
Cryo-EM structure of BIRC6 in complex with Smac. **a**, TR-FRET based displacement assay. Increasing concentrations of Smac, casp-7, HtrA2-S306A and casp-3 were titrated into BODIPY-labeled HtrA2-S306A at 50 nM, Flag-BIRC6 at 10 nM and Tb-anti-Flag antibody at 8 nM. IC50 values of 9±1 nM, 167±31 nM, 383±60 nM and 1358±148 nM were determined for Smac, casp-7, HtrA2-S306A and casp-3, respectively. **b**, TR-FRET equilibrium binding assay of BIRC6 BIR-domain mutant, ΔHelix variant, a double mutant and BODIPY-labeled Smac. The single mutants show comparable *K*_D_ of 26±2 nM and 30±5 nM for BIRmut and ΔHelix, respectively, while the double mutant has no detectable binding. Data in a, and b, is represented as mean ± SD (standard deviation) from n=3 technical replicates. **c**, Local resolution-filtered map highlighting the connectivity between Smac (grey) and the BIR domains of BIRC6 (in color), indicated by dashed circles. **d**, Side view showing locally refined Smac density in the central BIRC6 cavity. Density for the additional helices is colored light green as the β-X2 domain. **e**, Close-up highlighting how the Smac dimer is arching over the CBM domains. The connection to the IBM groove on the BIR domains is indicated with a dashed line, and the shown AVPI peptide is modeled after PDB 1G73. The attachment points of the two additional helices with the β-X2 domain are indicated with matching spheres and residue numbering. Domain labels with solid background belong to one protomer, domain labels with white background indicate domains of the second protomer.

To test whether Smac binding is solely governed by interactions of the Smac IBM motif and the BIR domains, we repeated the experiment with the BIRmut variant (C328S/C331S) of BIRC6 (**Fig. 3b**). Smac still bound with an apparent *K*_D_ of 26 ± 2 nM, confirming the existence of a previously proposed BIR-independent binding site for Smac on BIRC6^34^. To characterize the binding mode, we collected cryo-EM data for a stable BIRC6-Smac complex and obtained an anisotropic reconstruction at a nominal resolution of 2.5 Å (**Extended Data Fig. 4d and Extended Data Table 2**). The resulting maps revealed clear additional helical density in the central cavity (**Fig. 3c,d**). 3D classification and local refinements enabled unambiguous placement of Smac using a previously determined crystal structure^35^ (aa12-184, **Fig. 3d,e**). Due to limited map quality in this region, the Smac N-terminus in the IBM binding groove on the BIR domain was not resolved. A local resolution-filtered map, however, revealed a connection between the modeled Smac N-termini and the BIR domains (**Fig. 3c**), which is accounted for by unmodeled residues (**Fig. 3e**). Surprisingly, we also identified clear density for two additional helices on top of Smac that are not from Smac itself (**Fig. 3d**). While the quality of the map prevented unambiguous building of the helix, careful inspection of secondary structure predictions and unmodelled parts identified it as an extended helix contributed by the BIRC6 β- X2 domain (aa1616-1666), connected to the β-X2 domain through long disordered linkers of ∼34 and 105 aa. The two helices, one contributed from each protomer, hold Smac in place on top of the CBM domains (**Fig. 3e**). To ask whether these helices constitute the BIR-independent secondary binding site^34^, we measured the equilibrium binding constants of a deletion construct of BIRC6 (ΔHelix, aa1616-1666 removed) to be 30 ± 5 nM, comparable to the BIRmut construct (**Fig. 3b**). A double mutant (BIRmut/ΔHelix) had no observable affinity to Smac (**Fig. 3b**), consistent with BIR domain and helix constituting the two binding sites. In vitro ubiquitylation assays confirmed that all mutant proteins still exhibited auto-ubiquitylation activity (**Extended Data Fig. 5a**), and that there is a cumulative inhibition of activity towards Smac (**Extended Data Fig. 5b**). While performing competitive binding assays, we noticed that Smac, once bound to BIRC6 exhibited, very slow off rates compared to HtrA2 with virtually no displacement visible up to 22h while HtrA2 was readily displaced (**Extended Data Fig. 5c,d**). This near irreversible binding of Smac is in accordance with the structural arrangement and explains how Smac can expel bound caspases when released from mitochondria and thereby drive apoptosis. These findings are also consistent with our data showing that BIRC6 ubiquitylation of casp-3 and casp-7 is effectively inhibited by the presence of Smac (**Fig. 1d**).

### Caspases bind to the BIR domain and reside in the central cavity

BIRC6 does not contain multiple BIR domains or the linker region which was found essential for XIAP binding to casp-3 and casp-7^28, 36^. We thus set out to characterize the binding of these caspases to full-length BIRC6. To reconstitute a stable complex between BIRC6 and casp-3 or casp-7, we added the small molecule caspase inhibitor Z-VAD-FMK to prevent proteolysis of BIRC6. Reconstructions from datasets collected for BIRC6 bound to casp-3 or casp-7 revealed a similar overall architecture for BIRC6 (**Extended Data Fig. 6**). In both structures, the caspase occupies the central cavity established by the two arms of the arch and the two CBM domains in a flexible manner, which manifests in blurred density (**Fig. 4a and Extended Data Fig. 6**). Despite the lack of high-quality density for the caspase, we could place crystal structures of the corresponding dimeric casp-3^28^, constrained by the position of the N-termini and the BIR domains (**Fig. 4b-e**). Connecting density (**Fig. 4a**) between the caspase and the BIR domains together with our mutational data (**Fig. 1e**) confirms that the dimeric caspases are recruited and held in place by canonical BIR domain interactions with the IBM of the caspase (**Fig. 4e**). The overall positioning of the caspase is further stabilized by extended charge and shape complementarity between the CBM domains and the caspases (**Fig. 4c-e**). The active site of the caspase, which in previous structures of the XIAP BIR2 domain bound to caspase-3 is occupied by the linker leading to BIR2^28^, is pointed towards the CBM domains. In some clusters (clusters 2 and 3 in **Fig. 4a**) there appears to be a connection between the CBM and caspase density, which could potentially be accounted for by a positively charged loop of the CBM (aa3186- 3196) reaching into the negatively charged active site (**Fig. 4c-e**).

**Fig. 4.**
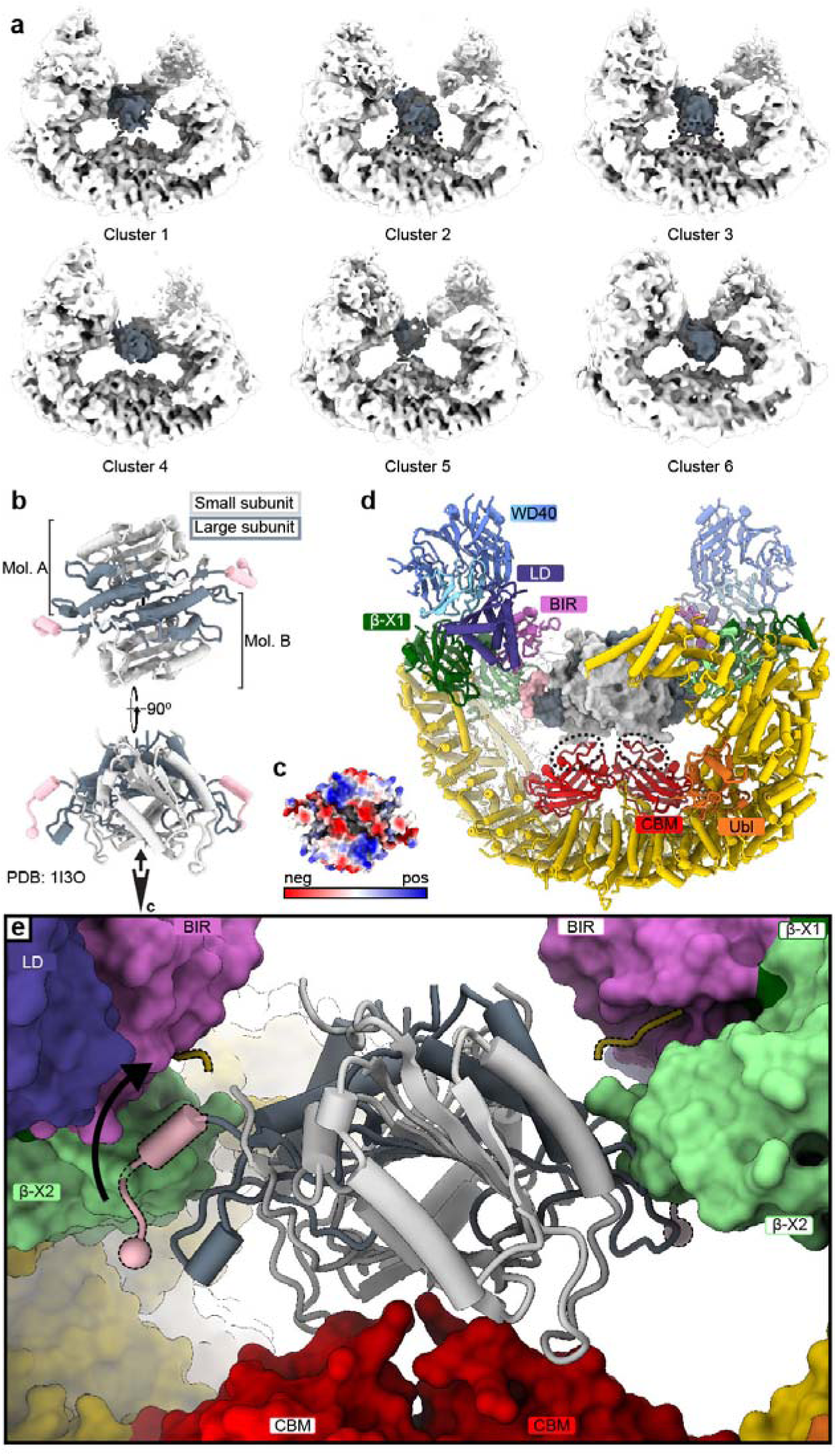
Cryo-EM structure of BIRC6 in complex with Casp-3. **a**, Six particle clusters after 3D variability analysis revealing a highly flexible binding mode of casp-3 in the central cavity of BIRC6. BIRC6 is shown in white, casp-3 in grey, and density connecting to the CBM domains in clusters 2 and 3 is indicated with dashed lines. **b**, Casp-3 structure from a published Casp-3/XIAP-BIR2 crystal structure (PDB 1I3O, BIR2 domain removed for clarity), with the large subunit in dark grey, the small subunit in light grey, and the IBM in salmon. Two orientations are shown and the viewing direction for panel c is indicated. **c**, Electrostatic potential map, viewed as indicated in b, revealing a negatively charged patch near the caspase active sites. **d**, Casp-3 shown as surface in same orientation as in (b, bottom), modeled in the central BIRC6 cavity indicating a snug fit. The positively charged loop (aa3186-3196) in the CBM is indicated by dashed lines. **e**, Zoomed-in view for the BIRC6/casp-3 model. BIRC6 is shown as surface, and the location of the IBM groove on the BIR domain is shown by a dashed yellow cartoon strand taken from PDB 1I3O. The IBM of casp-3 needs to swing up to reach into the BIR domain, the movement is indicated with a black arrow. Casp-3 shown as cartoon in same orientation as in (b, bottom) and (d).

### HtrA2 binds BIRC6 competitively with caspases and Smac

Similar to Smac, the serine protease HtrA2, which is primarily involved in clearance of misfolded proteins in the intermembrane space^37^, is also released from the mitochondria upon activation of the apoptotic cascade^38^. HtrA2 is a trimeric enzyme and after maturation in the mitochondrion consists of an N-terminal protease domain and a C-terminal PDZ domain, which is used for autoregulation and substrate recruitment^39–41^ (**Extended Data Figure 7a**). When released from the mitochondria, a canonical IBM (AVPS) is exposed, and HtrA2 has been shown to interact with BIRC6^19^. We sought to determine the structure of HtrA2 bound to BIRC6 to better understand how HtrA2 recognizes BIRC6. To prevent proteolysis, we used the active site mutant HtrA2-S306A, which was mixed with BIRC6, and a stable complex was purified and its consensus structure determined by cryo-EM at an overall resolution of 2.8 Å (**Extended Data Fig. 7b-e**). Density was observed in the central cavity of BIRC6 and, after 3D variability analysis followed by clustering, identified as a trimer of HtrA2-S306A (**Fig. 5a**). While making extensive contacts to the CBM domains and the helical arch structure of BIRC6, HtrA2-S306A is perfectly placed to be anchored by the BIR domains (**Fig. 5b,c**). HtrA2-S306A presents itself in a previously unobserved conformation that appears to be primed for activity (**Extended Data Fig. 7a**), with only one of the PDZ domains clearly defined by density (**Fig. 5a**). This intermediate conformation is stabilized by the Ubl domain in BIRC6 (**Fig. 5b,c**). Similar to Smac and caspases, HtrA2 interacts with the CBM domains primarily through an extended interface of charge complementarity (**Extended Data Fig. 7f)** and the overall conformation suggests that HtrA2, upon engagement with BIRC6, enters into a conformation that is no longer auto-inhibited explaining the observed proteolysis of BIRC6 by HtrA2 (**Fig. 1e**). In the observed arrangement, the active sites of HtrA2 are only ∼24 Å away from a potential HtrA2 recognition site (SYIF)^42^ which is located in a loop in the CBM domains (aa3198-3201, **Fig. 5c**), and the observed flexibility could well allow HtrA2 to cut BIRC6 at this site.

**Fig. 5.**
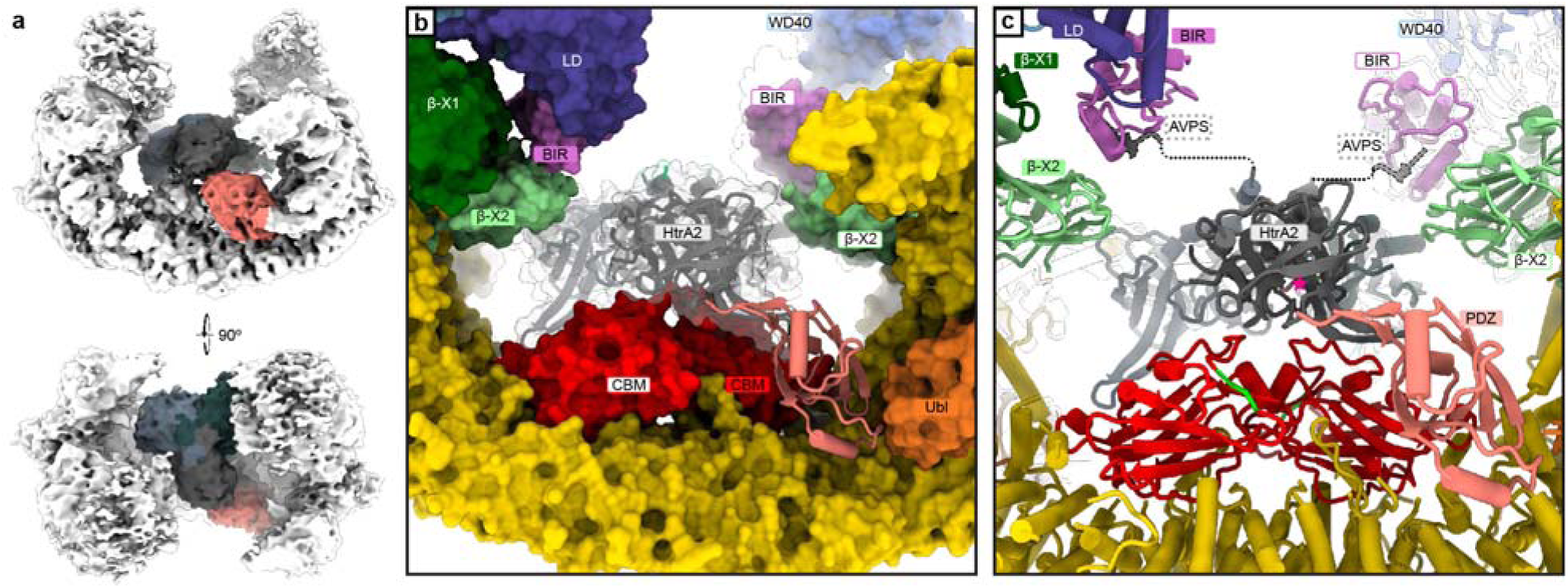
Cryo-EM structure of BIRC6 in complex with HtrA2. **a**, Cryo-EM density of BIRC6 (white) in complex with HtrA2-S306A (colored). Density was low pass-filtered to 5 Å. **b**, Detailed view with BIRC6 shown as surface representation and HtrA2-S306A as cartoon. The PDZ domain (salmon) wedges in between the Ubl and CBM domains, while the protease domains sit over the CBM domains. **c**, Detailed view with BIRC6 and HtrA2-S306A shown as cartoon. The location of the HtrA2-S306A active site closest to BIRC6 is indicated with a pink star, and the putative protease recognition site on BIRC6 (in the CBM) is highlighted green.

### Smac-mimetics have modest effect on BIRC6 activity

Given that BIRC6 is a negative regulator of apoptosis in cells^19^, and exhibits all the canonical IAP biochemical features, we asked whether Smac-mimetics designed to promote apoptosis in tumors by blocking caspase recruitment to IAPs would also inhibit BIRC6 activity. We obtained representatives of all classes of clinical stage Smac-mimetics: SM-406, Birinapant, GDC-0917, LCL-161, GDC-0152, ASTX660 (**Extended Data Fig. 7a**). To establish potency in vitro, we performed casp-3 ubiquitylation assays at increasing concentrations of each inhibitor (**Fig. 6a** and **Extended Data Fig. 7b**). Overall, we observe modest inhibitory activity, with a rank order of: LCL-161, GDC-0152, GDC-0917, SM-460 > ASTX660 > Birinapant. These findings are in line with the fact that these Smac mimetics were optimized for binding to cIAPs and XIAP^15^ and exhibit weak affinity for the BIR domain of BIRC6. Further, it is unlikely that bivalent Smac mimetics, such as Birinapant, can engage both BIR domains of BIRC6 simultaneously given the distance between the two. Hence, they will not exhibit improved efficacy, in line with the observed inability to effectively inhibit BIRC6-mediated ubiquitylation of casp-3 and casp-7.

**Fig. 6.**
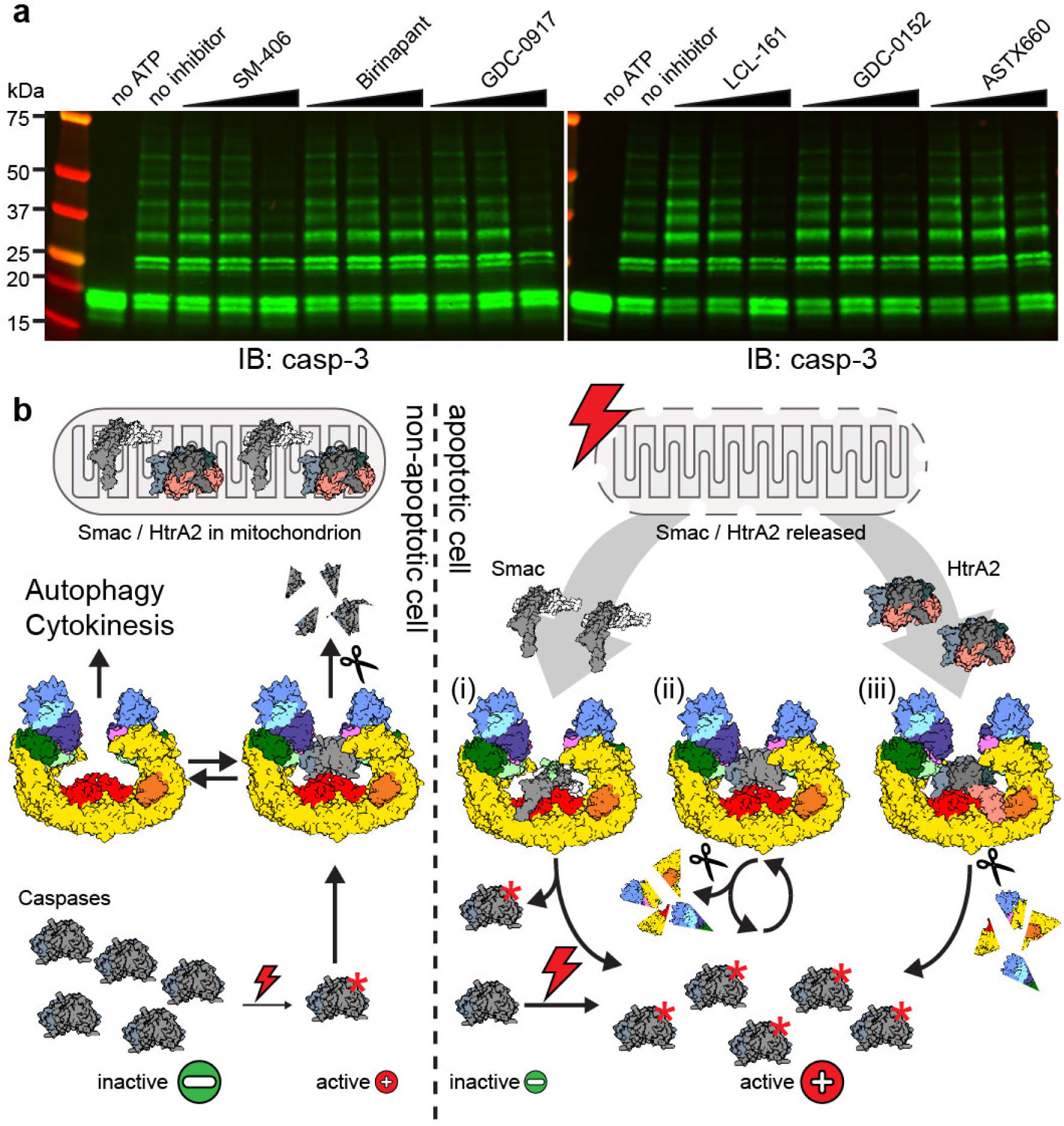
Smac-mimetics have limited effect on BIRC6 activity. **a**, In vitro ubiquitylation assays. In vitro ubiquitylation of casp-3 by BIRC6 in the presence of the indicated inhibitors was visualized by anti-casp-3 western blot. Blots are representative of three independent replicates. **b**, Cartoon summary illustrating the different regulatory networks in non-apoptotic (left) and apoptotic (right) cells. In non-apoptotic cells caspases are largely inactive and the small amount of active caspase is immediately cleared from the cell by proteasomal degradation. BIRC6 is able to regulate other pathways at the same time. Upon trigger of apoptosis Smac is released from the mitochondrion and the amount of active caspase is increased, leading to (i) inhibition of BIRC6- caspase binding by Smac and (ii) proteolytic cleavage of BIRC6 by active caspase (ii). HtrA2, while not able to efficiently free bound caspases, is also cleaving BIRC6 (iii), and all these mechanisms combined lead to a cumulative increase of active caspases.

## Discussion

While BIRC6 is an essential and highly evolutionarily conserved IAP^22^, a detailed understanding of its ability to regulate apoptosis through interactions with caspases, Smac and HtrA2 has been absent. Our structural and biochemical characterization of BIRC6 and its interaction with these key pro-apoptotic factors provides the molecular basis for BIRC6 activity as an IAP (**Fig. 6b**). The results presented here are consistent with previous cellular findings that BIRC6’s anti-apoptotic activity is dependent on the BIR and UBC domains^19^. We demonstrate that the dimeric architecture enables bivalent interactions of two BIR domains with multimeric substrates and regulators, and that it positions these in a central cavity (**Extended Data Movie 1**). This first example of a full-length IAP protein engaging its clients establishes a new paradigm for caspase recognition that does not involve distinct BIR domains and linker regions, but rather uses a single BIR domain and an extended central cavity to position caspases, Smac and HtrA2 in a mutually exclusive manner poised for regulation driven by differential affinities. This offers a solution to the long-standing conundrum of how Smac binding releases caspases and how single BIR domain containing proteins effectively bind to dimeric caspases.

BIRC6 serves multiple functions beyond regulating apoptosis, including mediating the late steps of cytokinesis^20, 23^ and negatively regulating autophagy^24, 25^ (**Fig. 6b**). Our structure reveals a large horseshoe-shaped scaffold built of helical repeats, with several functional domains present including an unpredicted WD40 propeller domain, a CBM-like domain and a ubiquitin-like domain. It is likely that these domains engage in protein-protein interactions relevant to roles in cytokinesis and autophagy. For example, binding of LC3B in the context of down-regulating autophagy has been mapped to the region between residues 2201 and 4300 on BIRC6^25^. This region encompasses the CBM-like and the Ubl domain, and future studies are needed to dissect the functional contributions of these individual domains. It is noteworthy that all functional domains appear arranged in a fashion that would potentially expose binding partners to ubiquitin ligase activity, especially considering the mobile nature of the UBC domain.

Our structures reveal significant differences in caspase binding to what had been observed in previous studies. Previously, casp-3 and casp-7 were found to interact with a linker region N-terminal to the BIR2 domain of XIAP, which binds in reverse over the active site of the caspase^28, 36^. In our structure of BIRC6, casp-3 and casp-7 seem to engage in a canonical BIR domain interaction with the N-terminal IBM directed towards the BIR domain. In addition, in our model casp-3 and casp-7 are stabilized by electrostatic interactions with the CBM domains confining them to the central cavity of BIRC6. This binding mode, together with the observation that Smac binding in the full-length BIRC6 context is nearly two orders of magnitude tighter with very slow off rates, explains how Smac binding is mutually exclusive and can thereby effectively release active caspases from BIRC6 (**Fig. 6b**). While additional studies are needed to see if similar principles may apply to other IAPs, our structures now provide a molecular understanding for Smac-mediated release of casp-3 and casp-7 from an intact IAP.

Drugs targeting anti-apoptotic pathways such as venetoclax are showing great clinical success and following the discovery and structural characterization of the Smac-IAP interaction, several groups developed Smac-mimetics to activate apoptosis for the treatment of cancer^15^. Despite several generations of Smac-mimetics with differing profiles in selectivity and binding modes, the clinical responses have been underwhelming^16^ (clinicaltrials.gov, 2022). All current Smac-mimetics were developed with a focus on XIAP and cIAPs as the role of BIRC6 and other family members as anti-apoptotic IAPs remained ambiguous. Our data demonstrates that clinically explored Smac mimetics are poor antagonists of BIRC6-mediated caspase inhibition and thereby spare this arm of IAP-mediated pro-survival signals. Our structures and biochemical characterization now offer ways to directly target BIRC6 through new Smac mimetics or by targeting the CBM domain of BIRC6.

Importantly, the mechanism observed for inhibition of caspases by BIRC6 and how it is counteracted by Smac provides insights into two distinct ways of inhibiting highly processive enzymes such as proteases or E3 ubiquitin ligases. While ubiquitylation and very tight, slow off-rate binding appear different at first, they both solve the problem of overcoming processivity through a kinetic component in which ubiquitin-mediated turnover leads to irreversible destruction of caspases and very tight binding of Smac to BIRC6 similarly leads to a near irreversible sequestration of BIRC6 allowing apoptosis to proceed beyond the point of no return. This elegant solution is likely more general in the regulation of such enzymes, and it is noteworthy that drugs targeting E3 ligases or proteases are also often characterized by very tight or irreversible binding.

While our study provides a comprehensive understanding of how the class II BIR domain protein BIRC6 interacts with casp-3, casp-7, Smac and HtrA2 and how these interactions allow for competitive regulation of activity, more work is needed to explore whether some of these observations also apply to class I BIR domain proteins such as XIAP or cIAPs. Our work offers a new mechanistic basis to dissect the individual contributions of IAPs and their interacting proteins lighting a path to new generations of apoptosis-inducing drugs.

## METHODS

### Cloning, protein expression and purification

A pDONR223-based entry clone (Thermo Fisher Scientific) containing the sequence encoding canonical full-length BIRC6 (aa1-4857, Uniprot: Q9NR09) was constructed from six gBlocks (IDT) in three consecutive rounds of Gibson assembly (NEB, E2621S). From there, the coding sequence was moved into a modified pDEST destination vector with an N-terminal FLAG tag and into modified pDARMO (pDarmo.CMVT_v1 was a gift from David Sabatini [Addgene plasmid #133072]) expression plasmids with N-terminal 3xFLAG- and Strep-tags. Mutant sequences were generated by excising pieces from these plasmids followed by replacement with gBlocks bearing the desired mutations using Gibson assembly. pcDNA3.1-HA-UBA6, containing the coding sequence for UBA6 (aa1-1052, Uniprot: A0AVT1) was a gift from Marcus Groettrup (Addgene plasmid #136995). UBA6 was subcloned into pDARMO with an N-terminal FLAG-tag. All BIRC6 variants and UBA6 were expressed transiently in Expi293 (Thermo Fisher Scientific, A14635) following the manufacturer’s manual. Cells were harvested 48-60 hours post transfection and lysed by sonication in lysis buffer (50 mM HEPES/KOH pH 7.4, 200 mM NaCl, 5% glycerol) supplemented with protease inhibitors. After clearance by ultracentrifugation (45 min, 120,000 g) the lysates were incubated with either FLAG-antibody-coated beads (Genscript, L004332-5) or Strep-Tactin®XT 4Flow high capacity resin (IBA life sciences, 2-5030-002). Bound proteins were eluted with 0.2 mg/mL 1xFLAG (DYKDDDK), 0.15 mg/mL 3xFLAG (MDYKDHDGDYKDHDIDYKDDDDK) peptide or 50 mM biotin, respectively. All FLAG-tagged variants were concentrated using centrifugal concentrators (Amicon, 30 kDa molecular weight cut-off (MWCO)) and polished by size exclusion chromatography (SEC, Superose 6 Increase, GE healthcare) in SEC1 buffer (30 mM HEPES/KOH, pH7.4, 150 mM NaCl, 3 mM Tris(2-carboxyethyl)phosphine (TCEP)). Strep-tagged protein was additionally purified by ion exchange chromatography: After elution from affinity resin, protein was loaded onto a Poros 50HQ column (Thermo Fisher Scientific, 1255911) and eluted with a linear NaCl-gradient from 200-750 mM. Elution fractions were concentrated and applied to SEC.

pET21b-Caspase-3-His_6_ and pET21b-Caspase-7-His_6_ were gifts from Clay Clark (Addgene plasmid #90087) and Guy Salvesen (Addgene plasmid #11825), respectively. The coding sequence for mature Smac without the N-terminal mitochondrial targeting sequence (aa56-239, Uniprot: Q9NR28) was ordered as gBlock and inserted into a pNIC-Bio2 (SGC Oxford) expression plasmid with a C-terminal His_9_ tag. A pET-20b-HrtA2(Δ133)-His_6_ expression plasmid was a gift from L. Miguel Martins (Addgene plasmid #14126). The MVPI/MVPS mutants for Smac and HtrA2 as well as the active site mutant (S306A) for HtrA2 were cloned by Q5 mutagenesis-style (NEB, E0554S) linearization with primers containing the desired mutations, followed by treatment with DpnI (NEB, R0176S), polynucleotide kinase (NEB, M0201S) and T4 DNA ligase (NEB, M0202S) for re-ligation. For expression, LBSTR *E. coli* expression strains (Kerafast, EC1002) were transformed, 2 L cultures were grown at 37 °C to OD_600_=∼0.6 and induced with 0.35 mM Isopropyl ß-D-1-thiogalactopyranoside (IPTG), except for casp-3 which was induced with 0.2 mM IPTG. Temperature was decreased to 18 °C, proteins were expressed overnight, and cell pellets were stored at −80 °C until processing. Cell pellets were resuspended in lysis buffer (50 mM HEPES/KOH pH 7.4, 200 mM NaCl, 20 mM imidazole, 5% glycerol, 1 mM TCEP) and lysed using sonication, followed by clearance using ultracentrifugation (60 min, 120,000 g). Cleared lysates were applied to high affinity Ni-charged resin (Genscript, L00223) and eluted with increasing imidazole concentrations (150 – 750 mM).

For casp-3, casp-7 and HtrA2-variants, elution fractions were concentrated using 3 kDa MWCO, 10 kDa MWCO and 30 kDa MWCO centrifugal concentrators, respectively, prior to polishing by SEC (Superdex75, GE Healthcare) in SEC2 buffer (30 mM HEPES/KOH, pH7.4, 200 mM NaCl, 3 mM TCEP). Smac variants were additionally purified using ion exchange chromatography: After elution from Ni-charged resin, elution fractions were diluted to ∼50 mM NaCl, applied to a Poros 50HQ and eluted with a linear salt gradient from 50-750 mM NaCl. Peak fractions were concentrated using centrifugal concentrators (10 kDa MWCO) and applied to a Superdex75 equilibrated in SEC2 buffer. Final protein samples of all constructs were flash-frozen in liquid nitrogen.

### In vitro ubiquitylation assays

In vitro ubiquitylation assays were carried out in a total volume of 15 µL with UBA6 and BIRC6 at 0.2 µM each, Mg-ATP (R&D systems, B20) at 1 mM, ubiquitin (R&D systems, U-100H) at 50 µM, and buffered with 1X E3 Ligase Reaction Buffer (R&D systems, B-71). Substrates and Z-VAD-FMK (UBPBio, F7111) were added at 6 µM and 50 µM, respectively. Caspase and inhibitor were pre-incubated for 30 min before mixing with BIRC6. If needed, BIRC6 variants were pre-incubated with Smac for 30 minutes at room temperature. Commercial BIRC6 inhibitors were added at 1 µM, 10 µM and 50 µM. Reactions were started with addition of ATP and run at 37 °C for one hour. Samples were separated on NuPAGE 3-8% Tris-Acetate (Thermo Fisher Scientific, EA0375BOX) or 4-20% TGX (Bio-Rad, 4561096) for visualization of auto-ubiquitylation or substrate ubiquitylation, respectively. For western blot analysis proteins were transferred onto PVDF membranes using an iBlot 2 dry blotting system (Thermo Fisher Scientific, IB21001). Specific primary antibodies were used to detect ubiquitin (Sigma-Aldrich, U5379), casp-3 (Cell Signaling, 9662S), casp-7 (Cell Signaling, 9492S), HtrA2 (Cell Signaling, 2176S) and Smac (Fisher Scientific, AF789) and blots were imaged using a LI-COR Odyssey CLx detecting an anti-rabbit secondary antibody (LI-COR, 926-32213). All assays were repeated at least three times.

### Stability assays

BIRC6 variants (1.85 µg) were incubated with casp-3 (0.46 µg), casp-7 (0.5 µg) or HtrA2 (2.85 µg) in a total volume of 20 µL in presence or absence of Smac variants (1.55 µg). Smac- containing reactions were pre-incubated for 30 min before addition of protease. Reactions were incubated for 20 min and 300 min for caspases and HtrA2, respectively. Reactions were stopped by addition of SDS-containing sample buffer and analyzed by SDS-PAGE using NuPAGE 3-8% Tris-Acetate gels. Assays were repeated at least three times.

### EM sample preparation and data collection

All data sets were collected using SerialEM^43^ (v3.8.0 for data set 1, v3.8.5 for data sets 2, 4 and 5, v3.8.6 for data set 3,) in a Thermo Scientific Titan Krios equipped with a Gatan Quantum image filter (20 eV slit width) and a post-GIF Gatan K3 direct electron detector. Movies were acquired at 300 kV at a nominal magnification of 105,000 x in counting mode with a pixel size of 0.825 Å/pixel for data sets 1, 2, 4 and 5, and 0.83 Å/pixel for data set 3. All grids (glow discharged Quantifoil 1.2/1.3 300) were vitrified in a Leica EM-GP operated at 10 °C and 90% relative humidity with 10 s pre-blot time.

*For data set 1 (BIRC6):* 4 µL FLAG-BIRC6 (2 mg/mL) with 0.8 mM CHAPSO were applied to grids and excess sample was blotted away for 3 s (3 s post blot). Three movies were recorded per hole with nine holes per stage position (19,647 movies total), resulting in 27 image acquisition groups. 50 frames were recorded per movie (2.4 s exposure time) with defocus varying from −0.8 – −2.0 µm with a total dose of 52.5 e^-^/Å^2^.

*For data set 2 (BIRC6/Smac)*: A ∼5-fold molar excess of Smac-His_9_ was added to FLAG-BIRC6 after elution from FLAG-antibody-coated beads and incubated at room temperature for 30 min. Imidazole was added to 30 mM and Smac-bound complex was enriched on Ni-charged resin. After elution with buffer containing 300 mM imidazole, sample was concentrated using centrifugal concentrators (30 kDa MWCO) and polished on a Superose 6 Increase equilibrated in SEC1 buffer. Final concentrated sample (1.4 mg/mL) was brought to 0.5 mM CHAPSO directly prior to grid preparation. 4 µL sample were applied twice to the grid and blotted for 3 s blot each time, followed by 3 s post-blot incubation before vitrification. 14,574 movies (50 frames each) were recorded with three exposures per hole and nine holes per stage position, resulting in 27 image acquisition groups. Defocus was varied from −0.8 – −2.2 µm, and total dose and exposure time were 56.4 e^-^/Å^2^ and 1.4 s, respectively.

*For data set 3 (BIRC6/casp-3):* Purified casp-3 (3.3 mg/mL) was incubated with 50 µM Z-VAD-FMK for 45 min at room temperature. Inhibited casp-3 was then added in a two-fold excess to Strep-BIRC6 (3.1 mg/mL) and incubated at room temperature for 15 min. Prior to grid preparation CHAPSO was added to 0.4 mM and 4 µL of BIRC6/casp-3 complex at 0.5 mg/mL were applied to grids, and then blotted for 3 s (5 s post-blot) before vitrification. Two movies (50 frames over 2.5 s, 10,422 movies total) were recorded per hole in nine holes per stage position (18 image acquisition groups), with a total dose of 51.4 e^-^/Å^2^ and defocus spreading from −0.8 – −2.0 µm.

*For data set 4 (BIRC6/casp-7)*: Casp-7 at 2.6 mg/mL was incubated with 50 µM Z-VAD-FMK for 45 min at room temperature and then added, in a two-fold molar excess, to Strep-BIRC6 at 3.1 mg/mL. After an additional incubation for 15 min at room temperature, CHAPSO was added to 0.4 mM immediately prior to grid preparation. 4 µl sample were applied and grids were blotted for 3 s with 2.5 s post-blot time before vitrification. Two movies with 51 frames each were recorded per hole (18,216 movies total), with 9 holes per stage position, resulting in 18 image acquisition groups. The total dose was 54.4 e^-^/Å^2^ spread over a exposure time of 3.1 s, with defocus values ranging from −0.8 – −1.9 µm.

*For data set 5 (BIRC6/HtrA2-S306A)*: A ∼4-fold molar excess of HtrA2-S306A-His_6_ was added to 3xFLAG-BIRC6 eluted from FLAG-antibody-coated beads and incubated for 30 min at room temperature. The complex was loaded onto a Superose 6 Increase SEC column equilibrated in SEC1 buffer. The final sample was concentrated to 1 mg/mL, brought to 0.4 mM CHAPSO and 4 µL were loaded onto grids before blotting for 3 s (1 s post-blot). 15,309 movies were recorded with 50 frames each. Three movies were recorded per hole with nine holes per stage position with defocus set to −0.8 – −2.2 µm. Total dose was 50.7 e^-^/Å^2^ and exposure time was 2.5 s.

### Data processing and model building

Relion-3.1^44, 45^, Relion-4^46^ as well as cryoSPARCv3^47^ and cryoSPRACv3.3.2 were used for all processing steps. All resolutions given are based on the Fourier shell correlation (FSC) 0.143 threshold criterion^48, 49^.

*Data set 1 (BIRC6):* 19,647 movies were corrected for beam-induced motion using UCSF MotionCor2^50^ and contrast transfer function (CTF) was estimated using CTFFIND4.1^51^. Poor quality micrographs (CTF fit >5.5 Å) were discarded. 4,058,599 particles were picked using crYOLO^52^. Standard clean up by 2D classification and homogeneous refinement in cryoSPARC led to an initial anisotropic reconstruction at 2.81 Å resolution from 823,642 particles. This reconstruction was used as seed class (six times) for two rounds of heterogeneous refinement of all initially picked particles. Particles (1,116,481) contributing to the resulting consensus refine were re-extracted in Relion (at 0.88 Å/pixel) and subjected to a total of three rounds of CTF-Refinement^53^ and one round of Bayesian polishing^54^. After re-importing into cryoSPARC, these polished particles led a consensus reconstruction at 2.07 Å (EMD-27832). For local refinement of the helical arch region, a soft mask was applied and the signal for the more flexible N- and C-terminal arm regions was subtracted. Local refinement led to a reconstruction at 1.98 Å (EMD- 27833). For local refinements of the N- and C-terminal portions, the consensus particle stack was symmetry expanded (C2), soft masks were applied and signal for other portions of the proteins was subtracted. For the N-terminal arm, the particles went through two rounds of classification by 3D variability^55^ followed by clustering with 16 classes and 8 classes, respectively. This led to reconstructions at 2.47 Å and 2.99 Å for the complete N-terminal arm (472,306 particles, EMD- 27834) and the very N-terminus (aa68-966, 325,666 particles, EMD-27835). The complete C2 symmetry expanded particle stack was used to calculate the map for the C-terminal arm (2,232,962 particles, 2.30 Å, EMD-27836). All final maps as well as maps post-processed using deepEMhancer^56^ were used for de novo model building in COOT^57^ (v0.9.8). All models were protonated (phenix.reduce) and refined against the deposited main maps (from cryoSPARC) using phenix.real_space_refine^58, 59^(v.1.20.1-4487). For EMD-27835 a model for aa1-1000 was generated by alphafold^60^, low confidence regions were pruned, and the individual domains were docked as rigid bodies. This initial fit was improved by iterative cycles of refinements in ISOLDE^61^ (v1.3) and phenix.real_space_refine and manual inspection and building in COOT.

All resulting models were deposited in the PDB under accession codes 8E2D (consensus), 8E2E (helical arch), 8E2F (N-terminal arm), 8E2G (aa68-966 of N-terminal arm) and 8E2H (C-terminal arm). The individually deposited maps and models have overlapping regions and for visualization (**Fig. 2**) a composite map and model were generated from the individual maps in UCSF ChimeraX^62^. Data collection parameters and refinement statistics are available in **Extended Data Table 1**.

*Data set 2 (BIRC6/Smac):* 14,574 movies were corrected for beam-induced motion and CTF was estimated on-the-fly in cryoSPARC live and 3,192,460 particles were extracted after TOPAZ^63^ particle picking. After clean-up through two rounds of heterogeneous refinement in cryoSPARC, the resulting particles (1,832,031) were re-extracted in Relion (at 1.2 Å/pixel) from micrographs that were pre-processed by MotionCor2 and CTFFIND4.1 implemented in Relion. After three rounds of CTF-refinement and two rounds of Bayesian polishing, particles were re-imported into cryoSPARC and further classified by two rounds of 3D variability analysis (16 classes and 8 classes, respectively) using a soft mask around Smac. The resulting particles (192,025) led to a reconstruction at 3.04 Å (EMD-27837) and a locally refined reconstruction at 3.44 Å (EMD- 27838). The combined model from data set 1 was fit into the density by domain-wise rigid body docking in ChimeraX followed by relaxation into the density using ISOLDE. A high resolution crystal structure of Smac (PDB 1FEW^35^) could readily be placed in the extra density, as well as two poly-Ala helices. The resulting model was protonated and refined with target restraints using phenix.real_space_refine, and deposited in the PDB as 8E2I (full model) and 8E2J (Smac and additional helices in locally refined map). Data collection parameters and refinement statistics are available in **Extended Data Table 2**.

Data set 3 (BIRC6/casp-3) and data set 4 (BIRC6/casp-7): Data set 3 (10,422 movies) and data set 4 (18,216) were processed analogously to data set 2 (**Extended Data Figure 6**), except that crYOLO was used for particle picking. The resulting polished particle stacks (671,917 and 880,343 for BIRC6/casp-3 and BIRC6/casp-7, respectively) yielded consensus reconstructions at 2.82 Å (BIRC6/casp-3, EMD-27839) and 3.01 Å (BIRC6/casp-7, EMD-27840). 2 rounds of masked 3D variability clustering (8 clusters and 6 clusters for BIRC6/casp-3, 8 clusters and 10 clusters for BIRC6/casp-7) revealed additional density in the central cavity, but no classes could be found where density was defined enough for actual model building. We conclude that this is due to pronounced flexibility and mobility of the interaction between caspases and BIRC6. The maps of the resulting clusters were deposited as additional maps along with the map from the consensus refinement under the corresponding EMDB identifier. A plausible model used for BIRC6/casp-3 (**Fig. 4**) was constructed by placing a crystal structure of casp-3 (PDB 1I3O^28^) in the central cavity where the extra density was observed, constrained by the distance of the N-termini and the BIR domains. Data collection parameters are available in **Extended Data Table 2**.

Data set 5 (BIRC6/HtrA2): Similar to the other data sets, data set 5 was pre-processed on-the-fly with cryoSPARC live and 2,049,607 particles were picked using crYOLO from 15,309 movies. 106,956 edge picks were filtered out by 2D classification (**Extended Data Fig. 7**) and the particle stack was cleaned by heterogeneous refinement. Particles were re-extracted in Relion (at 1.2 Å/pixel) from micrographs that were pre-processed by MotionCor2 and CTFFIND4.1 implemented in Relion. After three rounds of CTF-refinement and two rounds of Bayesian polishing, particles were re-imported into cryoSPARC and classified by one round of masked 3D variability analysis and visualized as 16 clusters. Two clusters revealed HtrA2-S306A in the same conformation and together (73,712 particles) yielded a reconstruction at 3.21 Å (EMD- 27841). The combined model from data set 1 was fit into the density by domain-wise rigid body docking in ChimeraX. The HtrA2 model was constructed by rigid-body fitting the N-terminal domains (as a trimer) and a single PDZ domain from a crystal structure (5M3N^41^) into the cryo-EM density. The model was protonated, refined against the main map with target restraints using phenix.real_sparce_refine and deposited in the PDB with identifier 8E2K. Data collection parameters and refinement statistics are available in **Extended Data Table 2**.

Maps sharpened with automatically determined B-values (from cryoSPARC) were deposited as main maps in the EMDB along with masks used in refinement and classifications, and all maps that were used for figure preparations or additionally assisted in model building were deposited as additional maps under the corresponding EMDB identifier (**Extended Data Table 3**). The dimer interface area was calculated using PDBePisa^64^, structural similarity searches were conducted using PDBeFold^65^ and all figures of models or density were generated in ChimeraX or COOT. Structural biology applications used in this project were compiled and configured by SBGrid^66^.

### Fluorescent protein labeling for TR-FRET assays

FLAG M2 (Sigma-Aldrich, F1804)) antibody at a concentration of 2.4 µM in SEC buffer was labeled with a 4-fold molar excess of Terbium(Tb)-Pfp ester (NCP311-Tb)^67^ for 75 minutes at room temperature. The reaction was quenched by the addition of 100 mM Tris pH 8, incubated for 10 minutes, and buffer-exchanged using Zeba Spin Desalting Columns (Thermo Fisher Scientific, 89882) back into SEC2 buffer. The degree of labeling (DOL) was determined to be 2.57, using extinction coefficients of 272,440 M^-1^cm^-1^ for the antibody and 22,000 M^-1^cm^-1^ (at 340 nm wavelength) for the Tb-complex. For labeling of Smac, a single cysteine was introduced directly preceding the C-terminal His-tag, and the protein (Smac-Cys) was purified following the protocol established for wild-type Smac. Smac-Cys (149 µM), casp-3 (101 µM) and casp-3 (74 µM) were labeled with equal molar amounts (for Smac) or 2-fold molar excess (for caspases) of BODIPY FL Maleimide (Thermo Fisher Scientific, B10250) for 30 min (Smac) and 60 min (caspases) at room temperature. Reactions were quenched by addition of 10 mM 2- Mercaptoethanol and buffer exchanged using Zeba Spin Desalting Columns. HtrA2-S306A at 185 µM was labeled with a 2-fold molar excess of BODIPY FL NHS ester (Thermo Fisher Scientific, D2184) for 60 min at room temperature, quenched with 100 mM Tris pH 8, and excess dye was removed by buffer exchange into SEC2 buffer using Zeba Spin Desalting Columns. DOLs for BODIPY labeled proteins were determined to be 0.48, 0.58, 0.6 and 0.48 for Smac-Cys, casp-3, casp-7 and HtrA2-S306A, respectively, using an extinction coefficient of 80,000 M^-1^cm^-1^ for BODIPY at 503 nm.

### TR-FRET binding and displacement assays

For equilibrium binding assays, 2 nM and 10 nM 3xFLAG-BIRC6 variants were mixed with 8 nM Tb-FLAG-M2 antibody and incubated at room temperature for one hour. BIRC6 variants were mixed in 384 microwell plates (Corning, 4514) with the corresponding BODIPY-labeled substrates, ranging in concentrations from 0.008 nM to 4 µM. The final reaction volume was 15 µL in assay buffer (30 mM HEPES, pH 7.4, 200 mM NaCl, 1% BSA, 0.05% Tween-20). To ensure that equilibrium was reached, plates were read out several times over the next 1 to 2 days using a PHERAstar FX (BMG Labtech) plate reader. Tb was excited at 337 nm and emissions at 490 (Tb) and 520 (BODIPY) were recorded with a 70 µs delay over a window of 600 µs, and plates were measured for 6 cycles. For background subtractions the same dilutions series were run but BIRC6 was omitted, and the resulting values were subtracted from all measurements. The TR-FRET ratio of 520/490 was extracted for each data point and *K*_D_ values were fitted using the One-site (specific binding) equation in GraphPad Prism 9. To inhibit proteolysis, caspases were incubated with 50 µM Z-VAD-FMK for one hour before setting up the dilution series and Z-VAD-FMK concentration was held constant in all conditions that contained caspases. No substantial changes in *K*_D_ were observed for BIRC6 wild-type after 23 hours and for BIRC6 mutants after 28 hours of incubation. Interaction of BIRC6 wild-type with Smac and casp-7 was too tight to be measured accurately in our experimental setup, and for measurement of interaction with HtrA2-S306A we used the values derived from 10 nM BIRC6 series. Binding of casp-3 was too weak to be accurately determined (**Extended Data Fig. 4b,c**).

For displacement assays displacing BODIPY-HtrA2-S306A, 10 nM 3xFLAG-BIRC6 was incubated with 3.35 nM Tb-FLAG-M2 antibody and 50 nM BODIPY-HtrA2-S306A for 3 hours at room temperature. Mixtures with unlabeled proteins were set up as above and read out with the same settings several times over the following 1 to 2 days on a PHERAstar FX. No substantial changes were detected after 28 hours and the data were fitted using the [inhibitor] vs. response (four parameters) equation in Prims 9. All assays were performed as technical triplicates. For displacement assays displacing BODIPY-Smac, the reaction mixtures were done accordingly, but using 5 nM 3xFLAG-BIRC6 and 5 nM BODIPY-Smac.

### Data availability

Cryo-EM maps and coordinates have been deposited in the EMDB and PDB, respectively, under accession codes EMD-27832 (BIRC6 consensus, PDB: 8E2D), EMD-27833 (BIRC6 helical arch, PDB: 8E2E), EMD-27834 (BIRC6 N-terminal arm, PDB: 8E2F), EMD-27835 (BIRC6 N-terminal arm (aa68-966), PDB: 8E2G), EMD-27836 (BIRC6 C-terminal arm, PDB: 8E2H), EMD-27837 (BIRC6/Smac full, PDB: 8E2I), EMD-27838 (BIRC6/Smac local refine, PDB: 8E2J), EMD-27839 (BIRC6/casp-3 with clusters), EMD-27840 (BIRC6/casp-7 with clusters), EMD-27841 (BIRC6/HtrA2-S306A, PDB: 8E2K). Uncropped gels and western blot source data is available as **Supplementary Information 1**.

## Acknowledgments

This work was supported by the National Institutes of Health (NIH) grant NCI P01CA066996 and a Mark Foundation Emerging Leader Award 19-001-ELA (both to E.S.F.). We thank the staff at the Harvard Cryo-EM Center for Structural Biology for their outstanding support during grid screening and data collection. We acknowledge the SBGrid consortium for assistance with software and high-performance computing. We thank N. C. Payne and R. Mazitschek for providing the NCP311-Tb. We also thank Eric Bennett, Mike Eck, Nico Thomä, Wayne Fairbrother and members of the Fischer lab for valuable input and critical feedback on the manuscript.

## Author contributions

M.H. and E.S.F. conceived study and designed research plan. M.H. cloned and purified proteins and conducted all biochemical assays and cryo-EM structure determination. C.Y.J. performed activity assays with commercial inhibitors. M.H. and E.S.F. designed experiments and all authors analyzed and interpreted data. E.S.F. supervised the study and acquired funding. M.H. prepared figures and M.H. and E.S.F. wrote manuscript. All authors approved of the final version of the manuscript.

## Materials and Correspondence

Further information and requests for resources and reagents should be directed to and will be fulfilled by Eric Fischer.

---

**Extended Data Table 1.** Data collection and refinement statistics for data set 1.

|  |  |  |  |  |  |
| --- | --- | --- | --- | --- | --- |
|  | Data set 1 – BIRC6 |  |  |  |  |
| Microscope | Thermo Fisher Scientific Titan Krios |  |  |  |  |
| Voltage (kV) | 300 |  |  |  |  |
| Camera | Gatan K3 |  |  |  |  |
| Magnification | 105,000x |  |  |  |  |
| Pixel size (Å) | 0.825 |  |  |  |  |
| Total electron exposure (e<sup>-</sup>/Å<sup>2</sup>) | 52.5 |  |  |  |  |
| Number of frames (no.) | 50 |  |  |  |  |
| Defocus range (µm) | -0.8 - -2.0 |  |  |  |  |
| Data collection software | SerialEM 3.8.0 beta |  |  |  |  |
| Micrographs collected (no.) | 19,647 |  |  |  |  |
| Total extracted particles (no.) | 4,058,599 |  |  |  |  |
|  | Map 1 | Map 2 | Map 3 | Map 4 | Map 5 |
|  | Consensus | Helical arch | N-terminal arm | N-terminal arm (aa68-966) | C-terminal arm |
| EMDB accession code | **EMD-27832** | **EMD-27833** | **EMD-27834** | **EMD-27835** | **EMD-27836** |
| PDB accession code | **8E2D** | **8E2E** | **8E2F** | **8E2G** | **8E2H** |
| Final particles used (no.) | 1,116,481 | 1,116,480 | 472,306 | 325,666 | 2,232,961 |
| Map resolution (Å) | 2.07 | 1.98 | 2.47 | 2.99 | 2.30 |
| FSC threshold | 0.143 | 0.143 | 0.143 | 0.143 | 0.143 |
| Model composition |  |  |  |  |  |
| Non-hydrogen atoms | 30,898 | 13,301 | 6,098 | 4,032 | 5,132 |
| Protein residues | 3,936 | 1,654 | 771 | 523 | 656 |
| *B* factors (Å<sup>2</sup>) |  |  |  |  |  |
| Protein | 76.55 | 24.20 | 51.93 | 63.95 | 32.08 |
| Water | - | 25.71 | - | - | 25.89 |
| R.m.s. deviations |  |  |  |  |  |
| Bond lengths (Å) | 0.003 | 0.007 | 0.003 | 0.003 | 0.003 |
| Bond angles (°) | 0.562 | 0.734 | 0.619 | 0.603 | 0.604 |
| Validation |  |  |  |  |  |
| MolProbity Score | 1.36 | 1.91 | 1.66 | 2.29 | 1.38 |
| Clashscore | 6.5 | 9.86 | 6.66 | 12.10 | 6.85 |
| Rotamer outliers (%) | 0.85 | 1.55 | 1.44 | 2.2 | 0.52 |
| Ramachandran plot |  |  |  |  |  |
| Favored (%) | 98 | 96.3 | 97.04 | 93.36 | 98.89 |
| Allowed (%) | 2 | 3.7 | 2.96 | 6.64 | 1.11 |
| Disallowed (%) | 0 | 0 | 0 | 0 | 0 |

**Extended Data Table 2.**
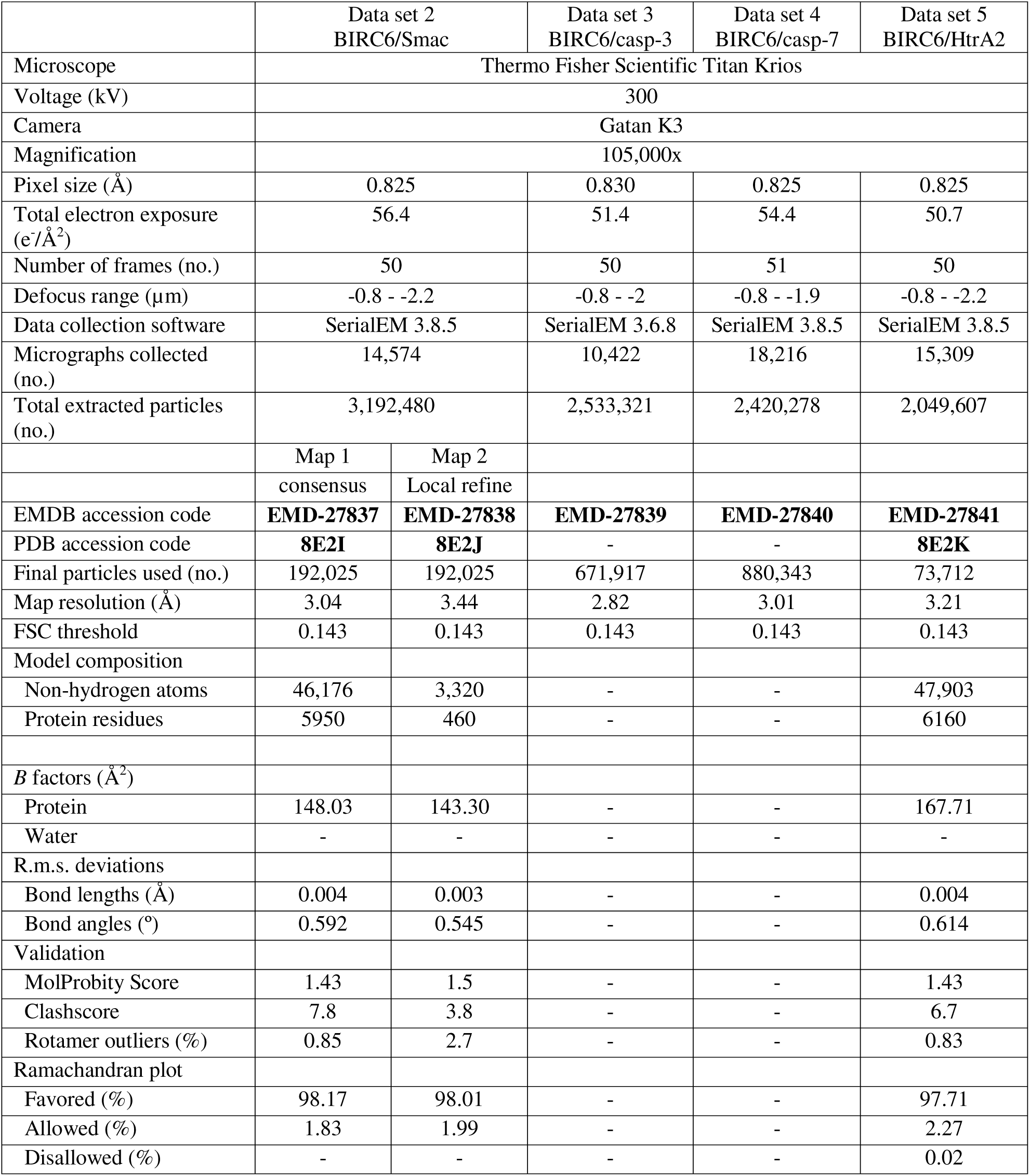
Data collection and refinement statistics for data set 2-5.

**Extended Data Table 3.**
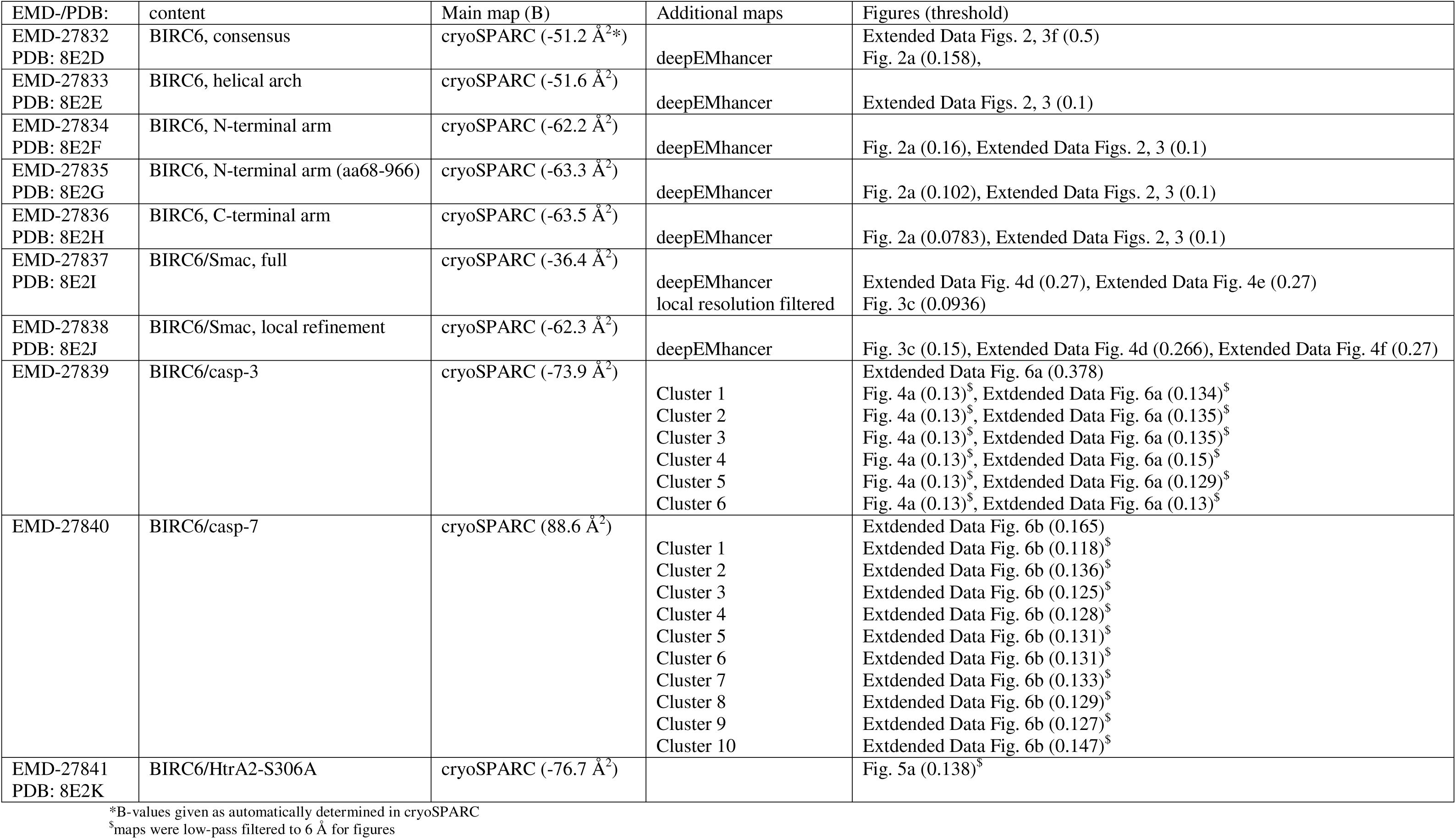
Overview of deposited maps, models, sharpening B-values and thresholds used in figures.

**Extended Data Fig. 1.**
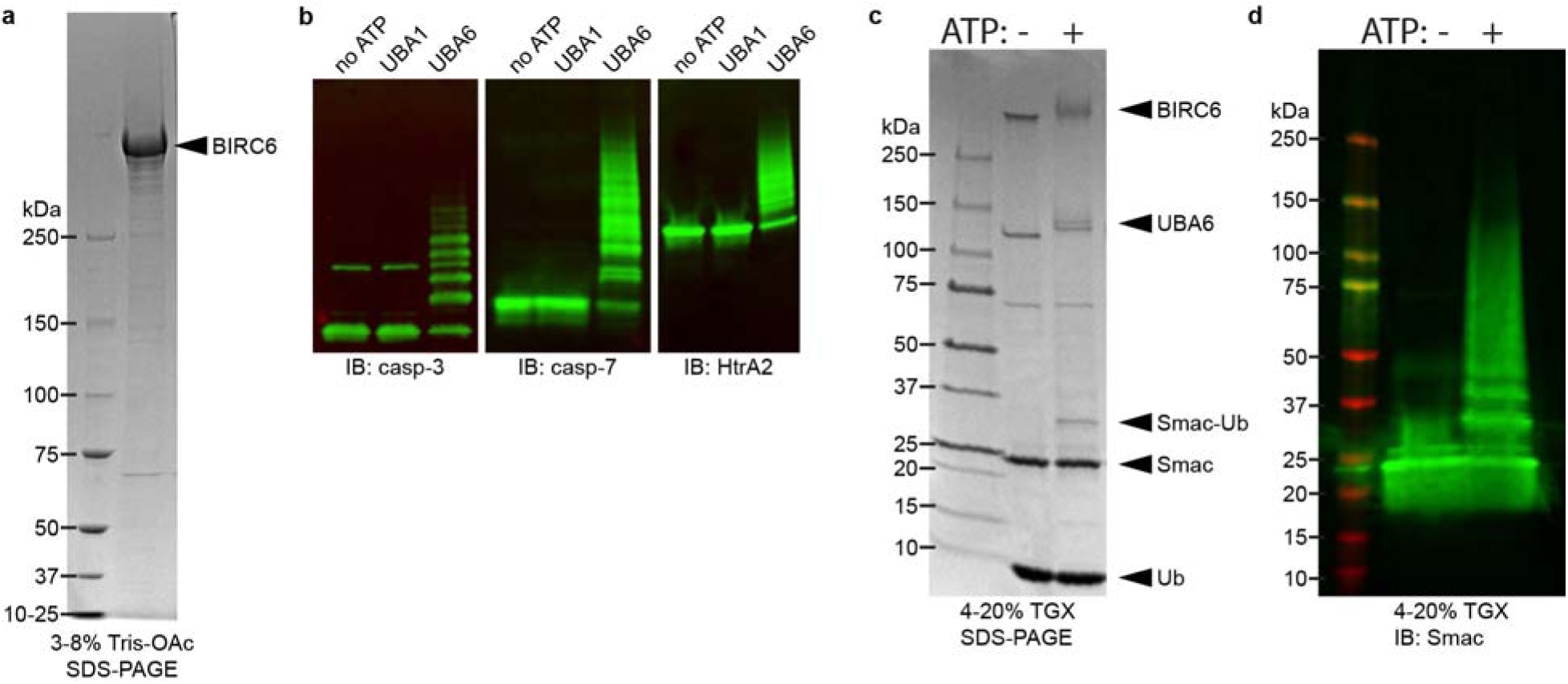
Purity and activity of BIRC6. **a**, SDS-PAGE analysis of final BIRC6 sample revealing highly purified protein with minimal degradation. **b**, In vitro ubiquitylation assays, analyzed by western blot (blotted for indicated substrates) using UBA1 and UBA6 as E1 enzymes. No activity can be observed when using UBA1. **c,d**, In vitro ubiquitylation assay using Smac as substrate, analyzed by SDS-PAGE (c) and western blot (d) blotted for Smac, establishing Smac as substrate. All gels are representative for at least 3 repetitions

**Extended Data Fig. 2.**
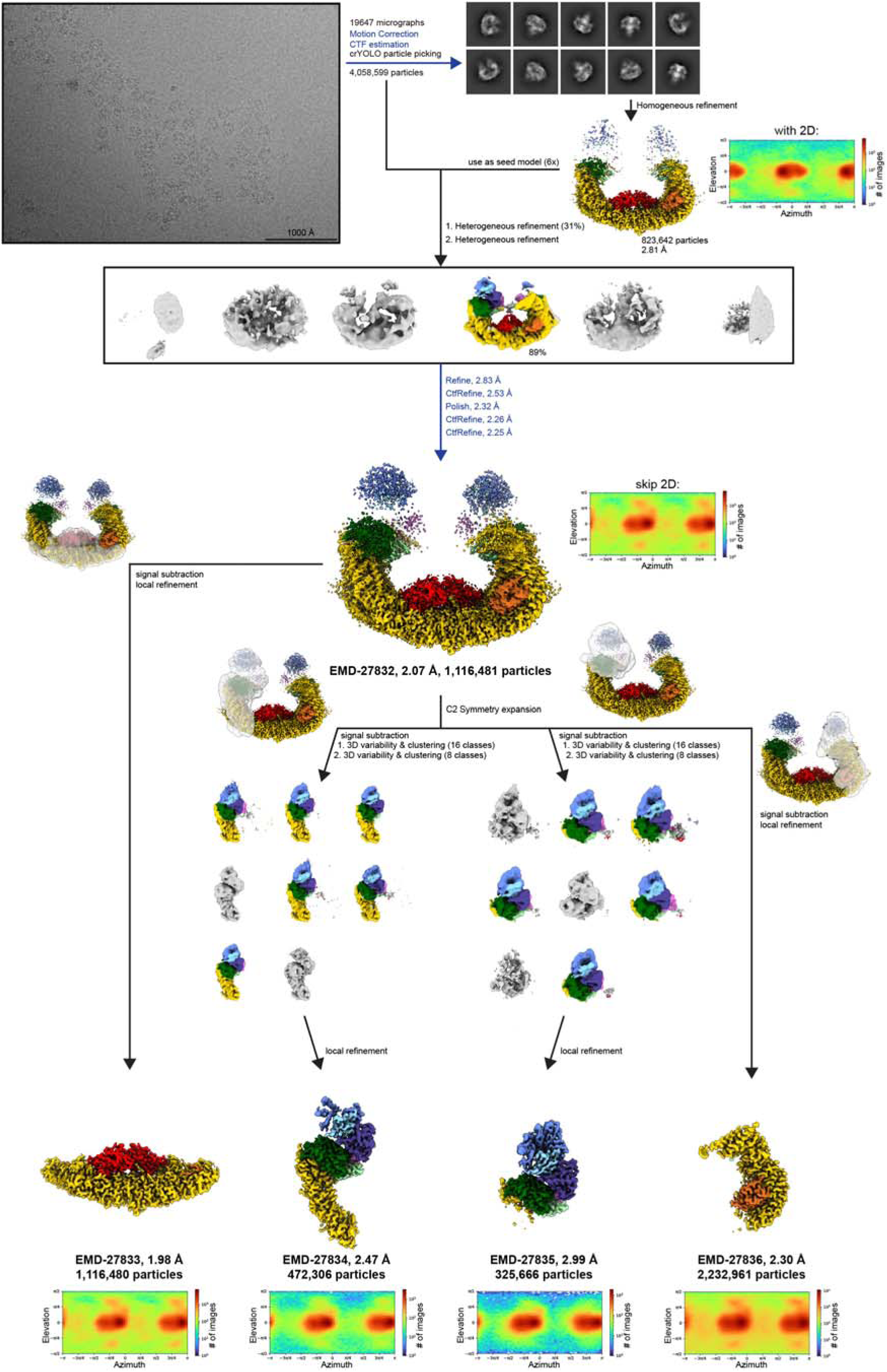
Cryo-EM processing workflow for the BIRC6 structure. **a**, Overview of processing workflow from raw micrograph (low pass-filtered to 10 Å, scale bar indicated) to final maps. Steps in blue were conducted in Relion, all others in cryoSPARC. All resolutions given are after post-processing. Particles belonging to colored volumes were taken into the subsequent steps. Contour levels of all final maps are available in **Extended Data Table 3**.

**Extended Data Fig. 3.**
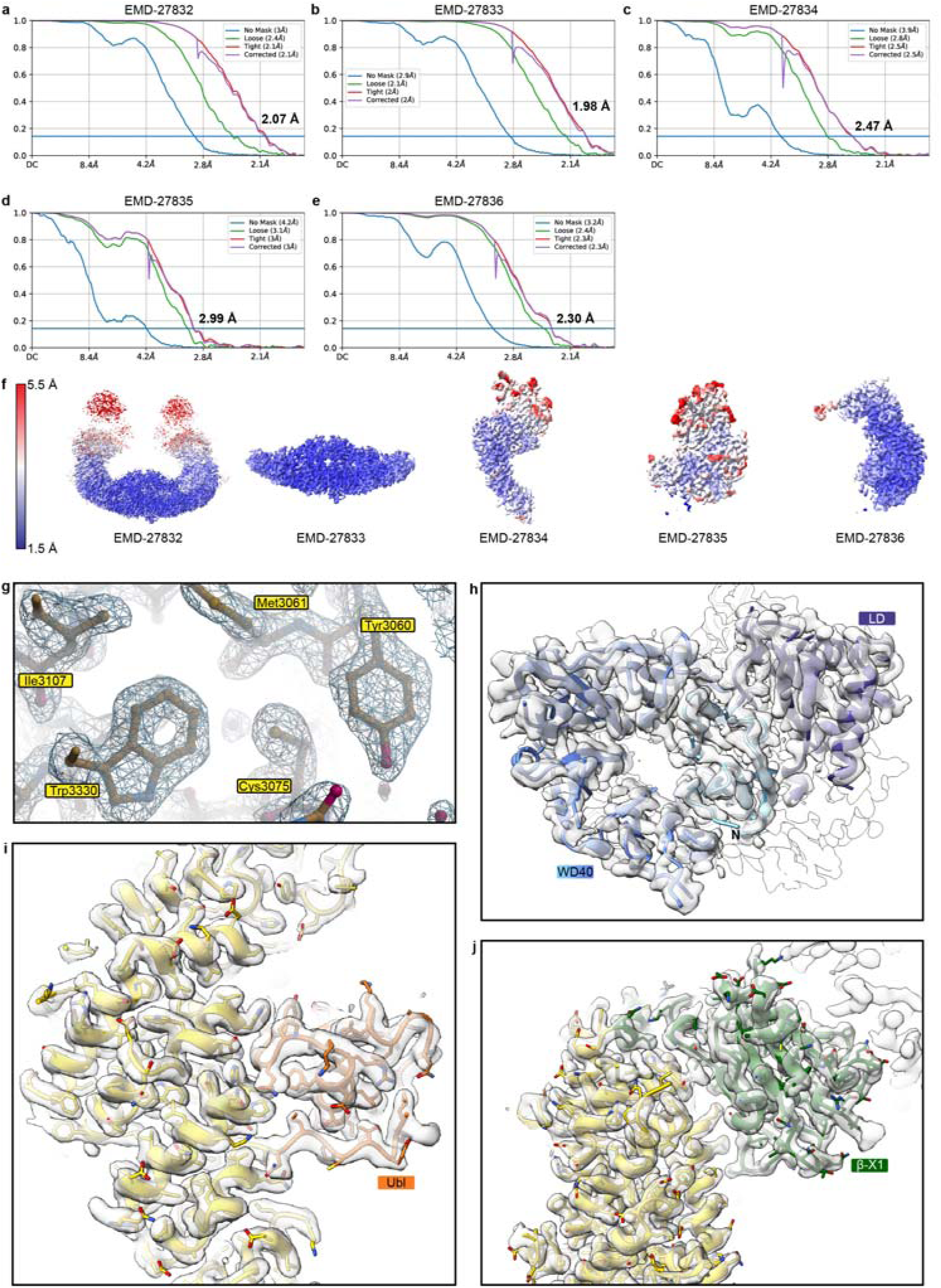
Map validation and quality for the BIRC6 structure. **a-e**, FSC plots for all deposited maps (EMD-27832 (a), EMD-27833 (b), EMD-27834 (c), EMD-27835 (d), EMD-27836 (e)). **f**, Final maps colored according to local resolution. **g-j**, Density examples for the helical arch region (g), the N-terminal WD40-like domain (h), the Ubl domain (i), and the β- X1 domain (j).

**Extended Data Fig. 4.**
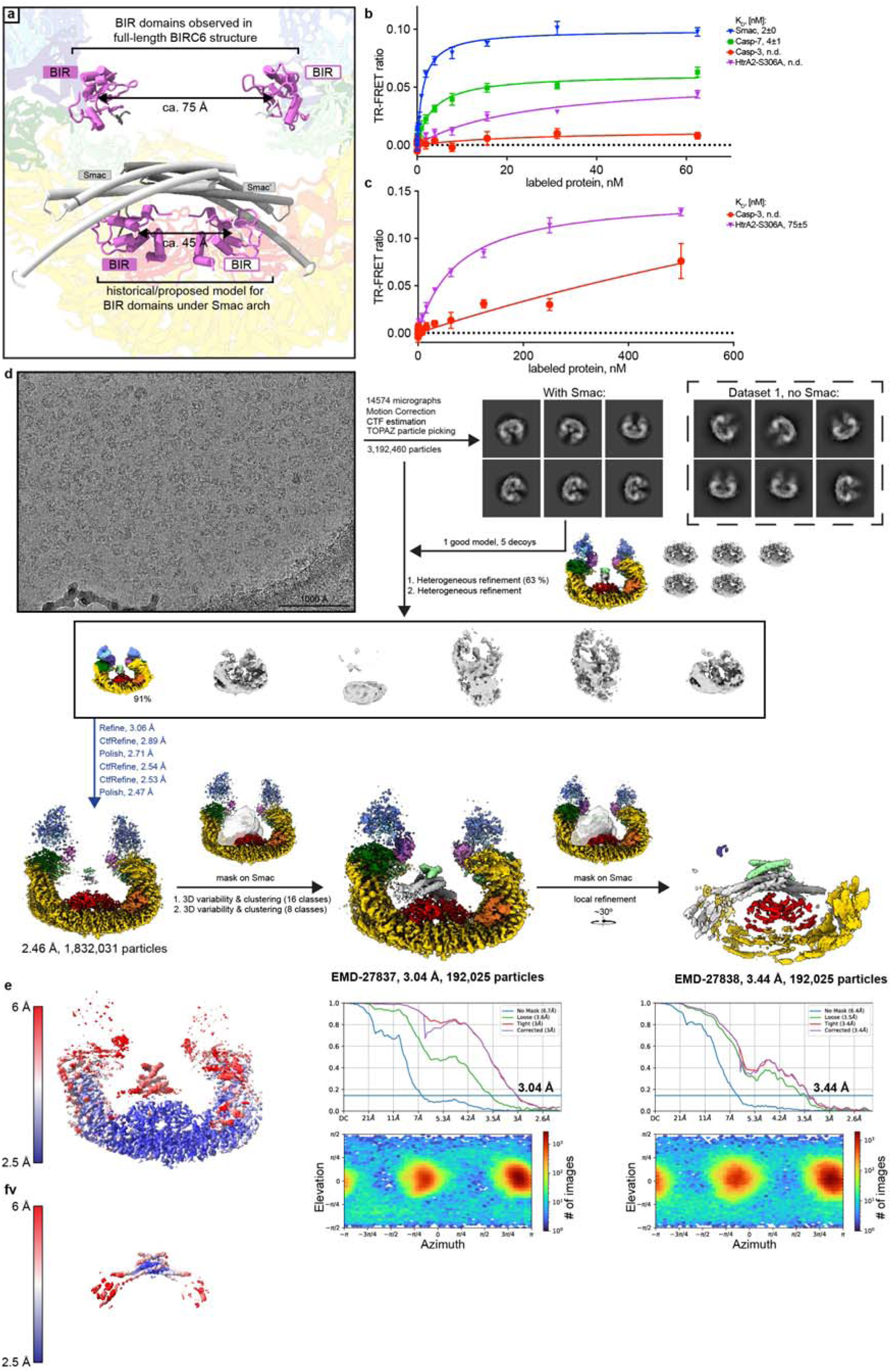
Cryo-EM processing workflow for the BIRC6/Smac structure. **a**, Comparison of previously proposed^29, 32, 33^ and observed binding arrangements of Smac and BIR domains. The distance in the model and observed in the presented structure are indicated. **b**, TR-FRET equilibrium binding assay of BIRC6 (2 nM) and BODIPY-labeled binding partners. Smac and casp-7 affinity exceed assay limitations (with *K*_D_ of 2±0 nM and 4±1 nM, rexpectively), while binding of casp-3 and HtrA2-S306A could not be determined. **c**, Same equilibrium binding titration as in b, but 10 nM BIRC6 was used. A binding constant for HtrA2-S306A (75±5 nM) could be determined. Data in b, and c, is represented as mean ± SD from n=3 technical replicates **d**, Overview of processing workflow for the BIRC6/Smac structure, from raw micrograph (low pass-filtered to 10 Å, scale bar indicated) to final maps. Marked in a dashed box are example 2D classes from data set 1, illustrating the differences in density in the central cavity. Steps in blue were conducted in Relion, all others in cryoSPARC. All resolutions given are after post-processing. Particles belonging to colored volumes were taken into the subsequent steps. Contour levels of all final maps are available in **Extended Data Table 3**. **e,f**, Local resolution mapped onto final maps.

**Extended Data Fig. 5.**
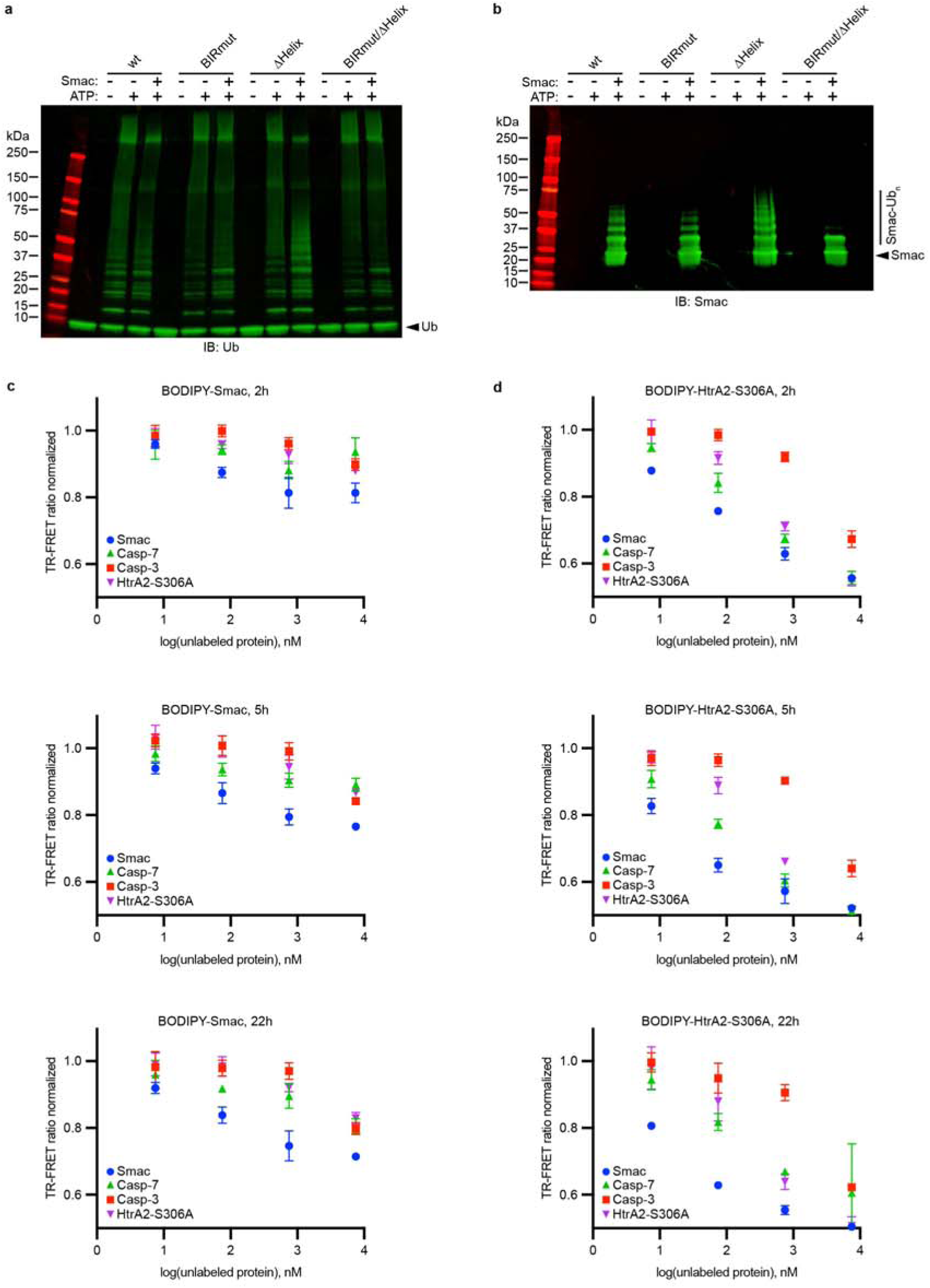
Activity of BIRC6 mutants and near-irreversible binding of Smac. **a**, In vitro auto-ubiquitylation assay. Western blot analysis (blotted for ubiquitin) reveals that all mutant proteins used in this study still exhibit normal auto-ubiquitylation activity. **b**, In vitro ubiquitylation assay, blotted for Smac, highlighting the cumulative inhibitory effect of the introduced mutations. a, and b, also highlight that auto-ubiquitylation activity is lower when Smac-directed activity is higher. Blots are representative for at least three independent replicates. **c,d**, TR-FRET displacement assays using BODIPY-Smac (c) and BODIPY-HtrA2-S306A (d) as tracer measured at various time point (2h, 5h, 22h). Over the course of 22h there is minimal displacement of BODIPY-Smac, but almost complete displacement of BODIPY-HtrA2-S306A. Data in c, and d, is represented as mean ± SD from n=3 technical replicates

**Extended Data Fig. 6.**
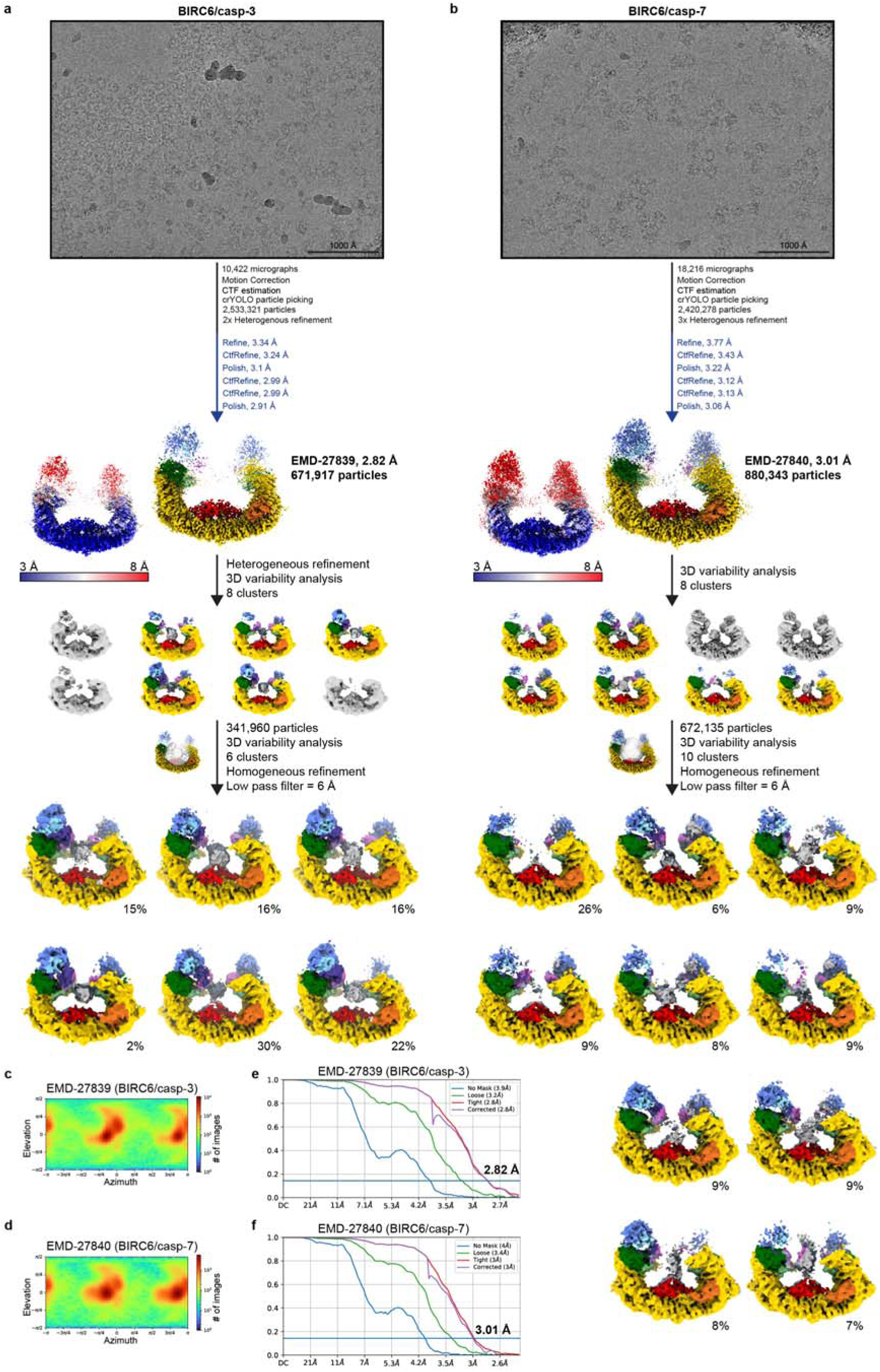
Cryo-EM processing workflow for the BIRC6-caspase complexes. **a,b**, Overview of processing workflow for the BIRC6/casp-3 (a) and BIRC6/casp-7 (b) datasets, from raw micrographs (low pass-filtered to 10 Å, scale bar indicated) to final maps. Steps in blue were conducted in Relion, all others in cryoSPARC. All resolutions given are after post-processing. Particles belonging to colored volumes were taken into the subsequent steps. Contour levels of all final maps are available in **Extended Data Table 3**. The maps corresponding to the final clusters were included as additional maps in the respective EMDB depositions. **c,d**, Viewing distributions plots for BIRC6/casp-3 (c) and BIRC6/casp-7 (d). **e,f**, FSC plots for the consensus refinements of BIRC6/casp-3 (e, EMD-27839) and BIRC/casp-7 (f, EMD-27840).

**Extended Data Fig. 7.**
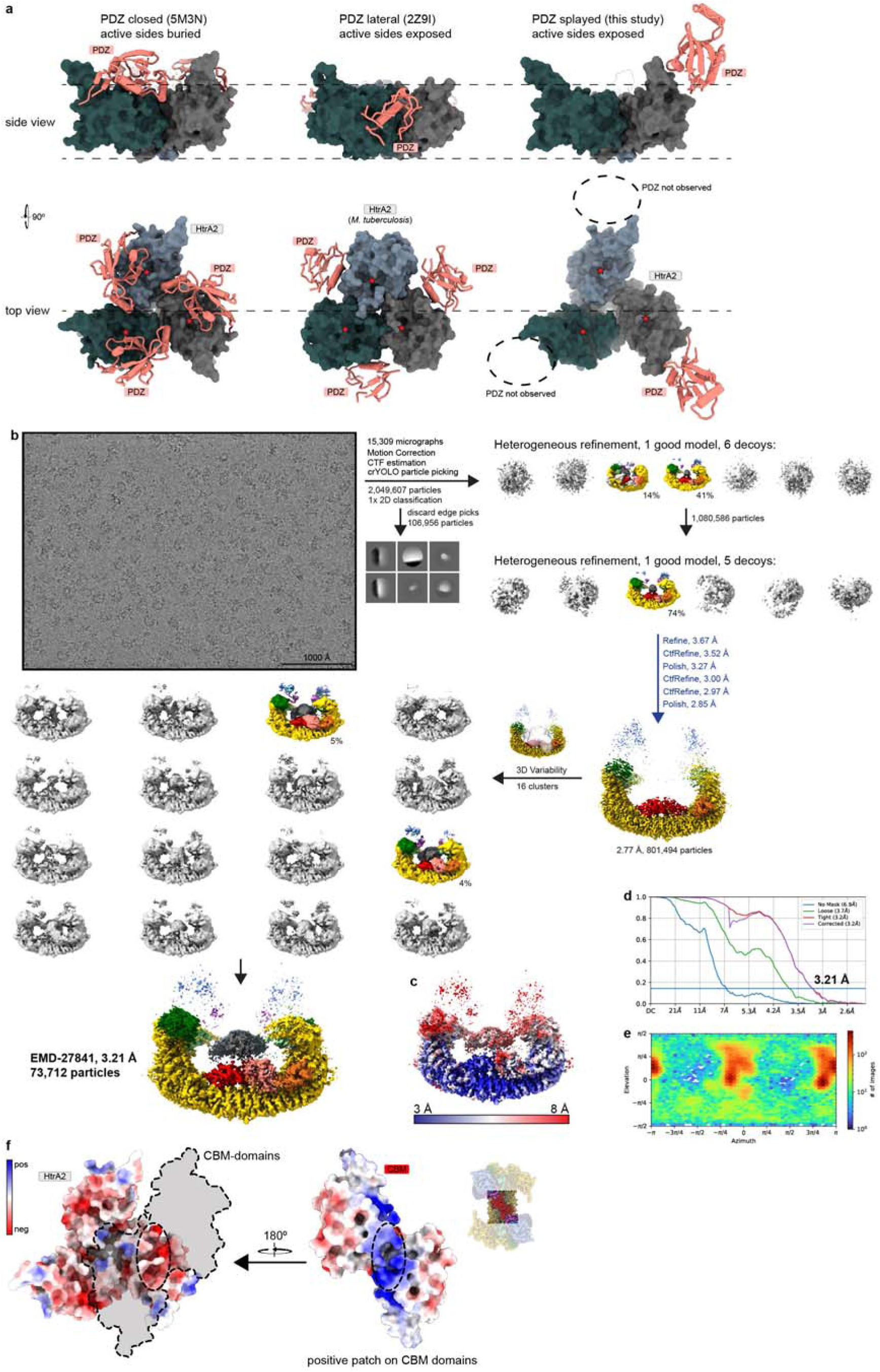
Cryo-EM processing workflow for the BIRC6/HtrA2-S306A structure. **a**, Three different conformations adopted by HtrA2. (left) Closed/inactive conformation observed in PDB 5MN3^41^. N-terminal protease domains are colored in shades of grey, and the PDZ domains, which occlude the active sites (marked with red star) are colored in salmon. Coloring is kept the same for all models and the plane established by the protease domains is indicated with dashed lines. (middle) Open/active conformation of *M. tuberculosis* HtrA2 (PDB 2Z9I^40^) with PDZ domains in the plane of the protease domains and accessible active sites. (right) Conformation as observed in this study, with exposed active sides and a PDZ domain in an intermediate conformation above the plane of the protease domains. **b**, Overview of processing workflow for the BIRC6/HtrA2-S306A structure, from raw micrograph (low pass-filtered to 10 Å, scale bar indicated) to final map. Steps in blue were conducted in Relion, all others in cryoSPARC. All resolutions given are after post-processing. Particles belonging to colored volumes were taken into the subsequent steps. **c**, Local resolution mapped onto final map. **d**, FSC plot. **e**, Viewing distribution plot. **f**, Coulombic surface coloring of Htra2 and the CBM domains, revealing an negatively charged patch over the active sites of HtrA2 which is directly covered by a positive patch found on the CBM domains.

**Extended Data Fig. 8.**
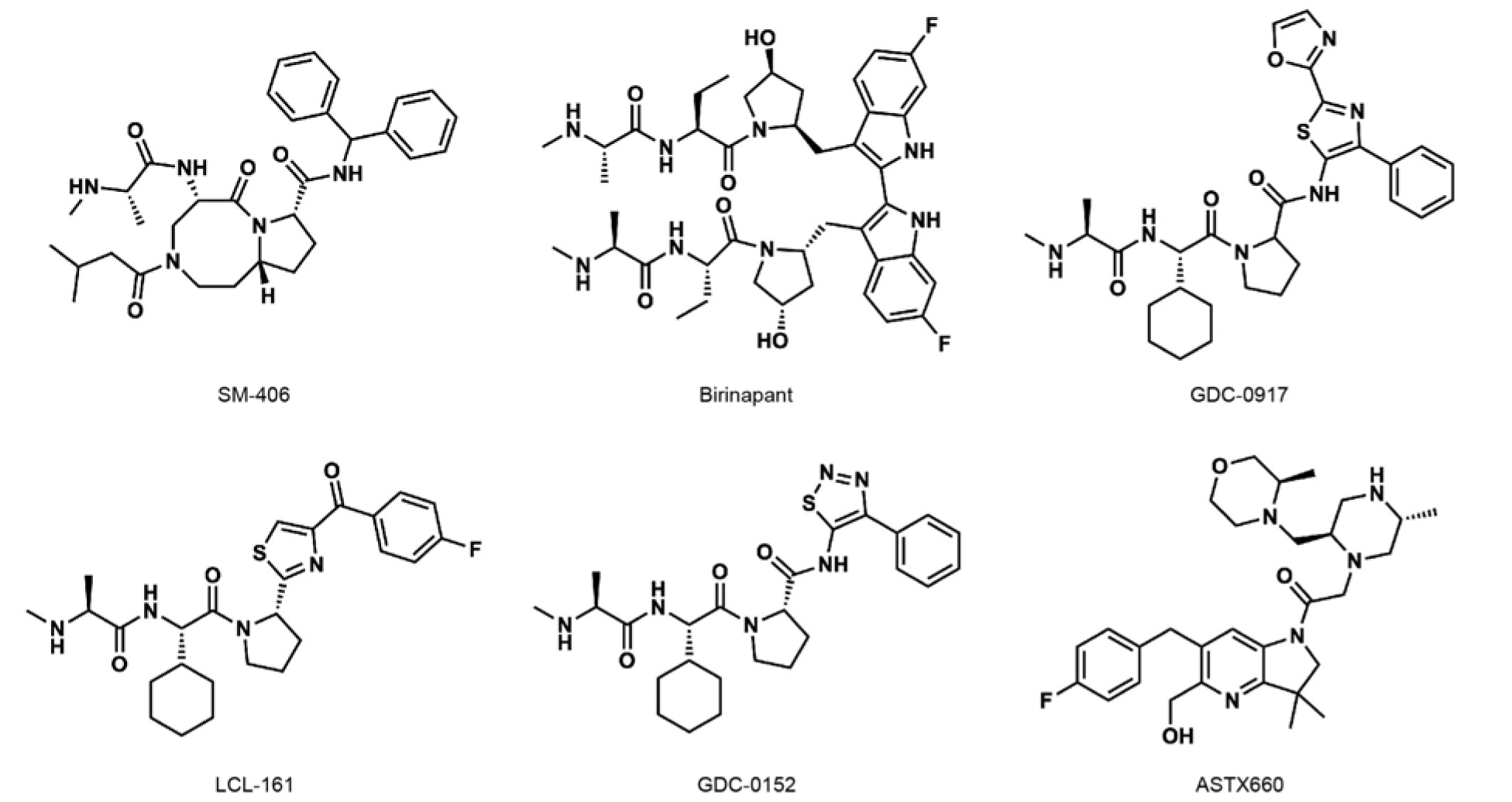
Commercially available IAP inhibitors. **a**, Chemical structures and names of commercially available IAP inhibitors used in this study.

**Extended Data Movie 1 | BIRC6 clients are flexibly bound in the central cavity.** Movie looping twice through manually aligned 2D classes (26 frames/classes each), illustrating the flexible manner in which BIRC6 clients are bound tin the central cavity.

**Supplementary Information 1 Uncropped gels and and western blots**

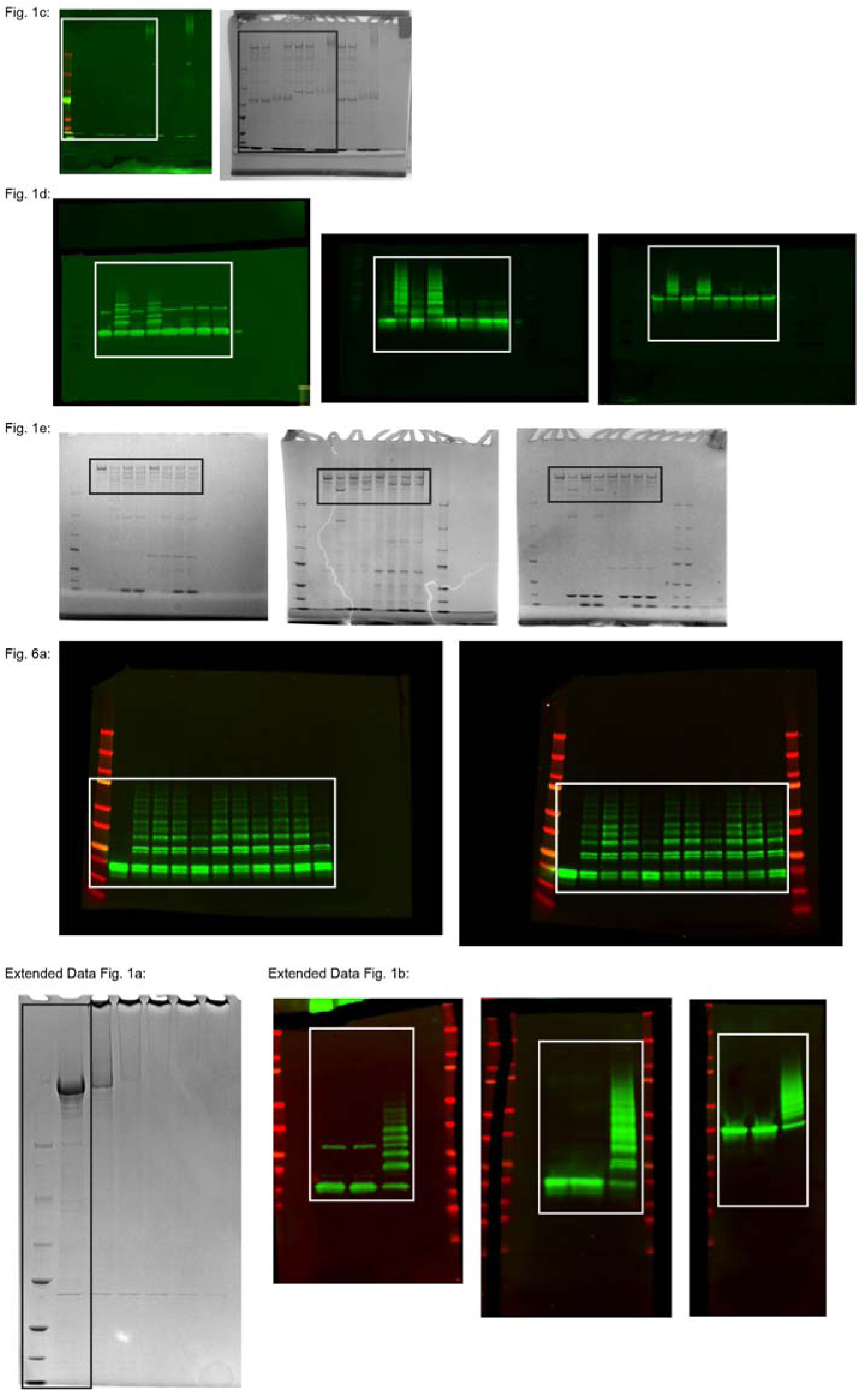
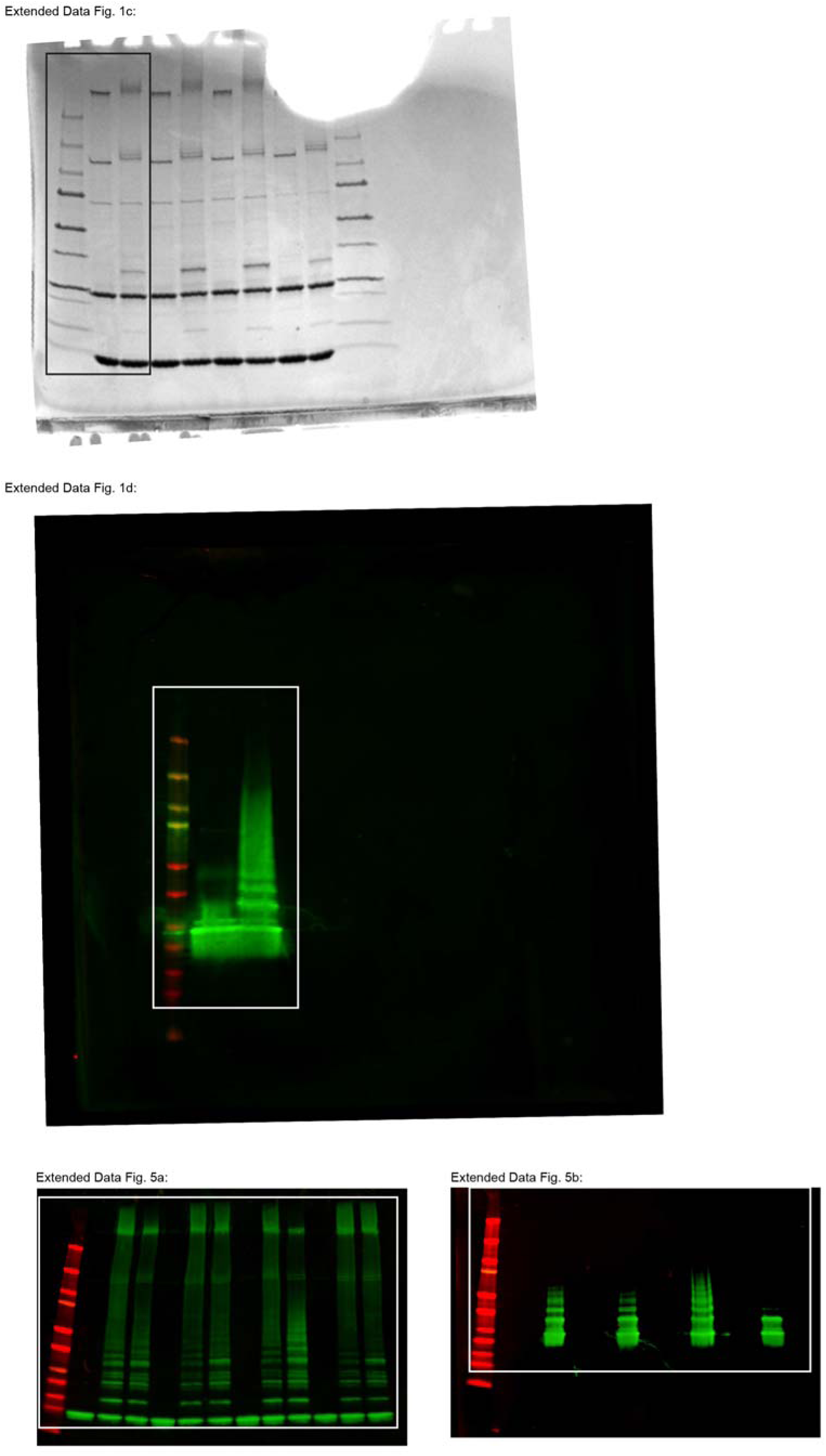

